# Fibrin barriers limit vancomycin penetration into staphylococcal communities and impact *S. aureus* transcriptional responses

**DOI:** 10.64898/2026.01.20.700504

**Authors:** Parsa Alba Farhang, Anjali Anil, Valerie Altouma, Timothy Huang, Rezia Era D. Braza, Kimberly M. Davis

## Abstract

During *Staphylococcus aureus* infection, bacteria are frequently organized into staphylococcal abscess communities (SACs), multicellular bacterial aggregates encased within a host-derived fibrin barrier. Fibrin barriers are thought to insulate SAC-resident bacteria from host immune cells, as strains unable to form these barriers exhibit marked attenuation of virulence in animal infection models. However, the clinical consequences of these structures on antibiotic penetration, and whether the structures also impact bacterial physiology and alter antibiotic susceptibility, remains understudied. Here, we built upon prior work to develop a three-dimensional (3D) collagen gel matrix-based model, utilizing human coagulation components, that supports high throughput *in vitro* staphylococcal community (SC) formation and is compatible with timelapse microscopy and bulk transcriptional profiling. Using this system, we show that the fibrin barrier protects SCs from vancomycin by restricting drug penetration. Furthermore, we show that the fibrin barrier impacts transcriptional responses, leading to both shared and distinct transcriptional profiles between WT and fibrin barrier-deficient SCs. Together, these findings expand our understanding of SC fibrin barriers beyond immune evasion to include modulation of antibiotic interactions and bacterial physiological state.

**AUTHOR SUMMARY:** Over the course of infection, many bacterial pathogens organize into clustered communities, which limits our ability to fully eliminate bacterial cells with antibiotic treatments. During *Staphylococcus aureus* infection, bacteria form dense clusters surrounded by a protective fibrin barrier derived from host blood clotting machinery, in structures known as staphylococcal abscess communities (SACs). These can be found in a range of different tissue sites, from the kidney, liver, and heart to the skin. SAC fibrin barriers have been shown to protect against immune cell invasion, but any effects on bacterial physiology and responses to antibiotics remain unclear. In this study, we developed an *in vitro* system to rapidly grow staphylococcal communities (SCs) in a 3D collagen gel matrix suspension, closely mimicking human tissue. Using this system, we show that fibrin barriers restrict vancomycin penetration into maturing SCs, protecting them from exposure to this clinically relevant antibiotic. These findings not only suggest that fibrin barriers serve as a physical barrier to drug diffusion but also show the importance of fibrin barriers in bacterial physiology and responses to vancomycin, highlighting their importance in SAC biology and infection persistence.

## INTRODUCTION

*Staphylococcus aureus* is a common causative agent of bacterial infections worldwide, resulting in a high degree of morbidity and mortality (1–5). Effective therapeutic options are scarce due to the rising prevalence of antibiotic resistance, as well as coordinated bacterial morphological structures that make infection clearance challenging (3,6–9). One of these structures is the staphylococcal abscess community (SAC), which is a tight cluster of bacteria encased in a fibrin pseudocapsule and more distal fibrin meshwork, derived from the host coagulation cascade, and surrounded by immune cells (8,10,11). Staphylococci that cannot form SACs, such as genetic mutants, are markedly attenuated in their virulence in mouse infection models (10,12,13). Additionally, there is preliminary evidence showing that SACs may protect against certain antibiotics, further highlighting their importance in clinical contexts (14). Expanding our understanding of how these structures form, are maintained, and are disrupted, will provide knowledge that can inform therapeutics aimed at improving clinical outcomes.

Specific virulence factors have been implicated in SAC development and maintenance. The accessory gene regulator (Agr) two-component system (TCS) is a quorum sensing system that controls the expression of a wide array of virulence factors through sensing of auto-inducing peptides (AIPs) released by individual bacteria (15–18). Several Agr-controlled virulence factors are implicated in SAC development, including protein A (*spa*), IsdA adhesin, and staphylokinase (8,10,17,19). Staphylokinase, a secreted protein that converts the clot disrupting enzyme, plasminogen, into its active form, has been shown to disrupt SC barrier integrity in a small subpopulation of SACs *in vitro*, possibly promoting bacterial dissemination (11,20). It has been shown that Agr is involved in SAC formation in a mouse skin infection model, but Agr is not required for kidney SAC formation (21,22). In addition to the Agr TCS, the *S. aureus* exoprotein (Sae) TCS also plays a key role in SAC biology. The SaeRS TCS, which responds to membrane perturbations, differential nutrient conditions, and oxidative stress, controls several virulence factors directly implicated in SAC formation, with coagulase (*coa*) being the most consequential (11,17,19,23–26). This TCS has been shown to be necessary for bacterial replication in skin and SAC formation in mouse kidney infection models (22,27,28).

Coa, and a distinct but functionally-related protein, von Willebrand factor-binding protein (vWbp), are enzymes that hijack the host coagulation cascade and help encase nascent SACs in their characteristic protective fibrin barrier (25,29). Although it is known that SaeRS promotes *coa* expression, it is not well understood how *vwb* expression is regulated, and this will not be a focus here (30). Both coagulase and vWbp individually promote blood coagulation through similar and somewhat redundant mechanisms (10,25,29,31). Each enzyme has an N-terminal region that binds and non-proteolytically activates the zymogen form of blood coagulation factor II (prothrombin) and a C-terminal region that binds the blood clotting glycoprotein fibrinogen (10,25,29,31,32). This fibrinogen-prothrombin complex converts soluble fibrinogen into insoluble fibrin that *S. aureus* uses to encase itself in physical fibrin barriers (11,12,25,29,31). Coagulase operates more proximally to the SAC, generating the fibrin pseudocapsule, while vWbp generates fibrin meshwork more distally (11). The resulting structure provides *S. aureus* with shelter that insulate bacteria from immune cells, making them important for *S. aureus* pathogenesis (8,11,12,14,19,31). Consistent with this, disruption of *coa* or *vwb* expression or protein function attenuates *S. aureus* virulence *in vivo* (10,11,18,20,31). However, fundamental questions remain regarding the role of fibrin barriers in SAC biology and pathogenesis in a clinical context. For example, only limited data exists to suggest a protective role of fibrin barriers against antibiotic penetration. It has been shown that gentamicin struggles to penetrate SACs and that exogenously disrupted SACs are more susceptible to gentamicin if treated at 100x the minimum inhibitory concentration (MIC), but assessing antibiotic susceptibility of SACs unable to form endogenous fibrin barriers with antibiotic concentrations close to the MIC, or mirroring serum circulating levels in patients, has yet to be done (11,12,14). Additionally, how SACs with and without fibrin barriers differ in development and transcriptional state, remains unknown. Addressing these gaps in knowledge could aid in improving antibiotic treatment outcomes.

Much of the work studying SAC formation has been accomplished by infecting mice with mutant *S. aureus* strains and performing endpoint analyses such as viability plating and histology. It is widely speculated that understanding when, in what host niches and in what bacterial subpopulations, the upstream virulence factors (Agr, Sae, Coa) are active is an important next step for understanding SAC biology and ultimately informing therapeutic approaches. Information gained from mouse models of infection has led to efforts to develop cell culture-based systems that recapitulate observations made in mice, that will enable further probing SAC biology, potential therapeutic interventions, and incorporation of human components (33–35). Traditional cell culture-based systems do not bridge this gap because they support growth in two dimensions (2D), failing to capture SAC spherical fibrin-encased morphology.

To address the limitation of traditional 2D cell culture-based systems, we have built upon recent work in other labs to develop an *in vitro* three-dimensional (3D) collagen gel matrix cell culture model to study staphylococcal communities (SCs) (11,14,36). This model involves seeding a bacterial-collagen gel matrix suspension into an individual well of a 96-well plate and then overlaying this matrix with media containing essential human coagulation components. Bacteria form fibrin-encased SCs in unique 3D planes, recapitulating key aspects of the structures found in mammalian infection (22). As shown here, host cells, bacterial fluorescent reporter strains, antibiotics, and bacterial mutants can be readily incorporated. Furthermore, each well in this 96-well plate-based system can contain its own matrix with its own unique microenvironment, making it a high throughput system to assess SAC biology. Another novelty of this iteration of the 3D collagen gel system is the ability to perform timelapse microscopy on fluorescent bacterial reporter strains as well as isolate bulk RNA for transcriptional profiling. Together, these features make this model high throughput, translationally relevant, and well positioned to capture nuances in SC biology at a high temporal resolution.

Here, we will use this model to answer questions regarding the role of fibrin barriers in SC biology and SC-antibiotic interactions: (i) whether fibrin barriers are protective against vancomycin and the potential protection mechanism, and (ii) whether the fibrin barrier or host blood components impact the phenotypic state of SC-resident bacteria and their ability to respond to vancomycin (37). Our results indicate that while fibrin barriers limit vancomycin penetration and maintain wild-type (WT) mature SCs in a transcriptionally distinct state relative to fibrin deficient mutants, they do not impact the ability of SC-resident bacteria to respond to vancomycin. Instead, it appears that both human plasma and human serum enable stationary phase bacterial populations to respond to vancomycin stress, something that we did not observe in standard laboratory media. Ultimately, this study further highlights the crucial role of fibrin barriers in SC antibiotic susceptibility as well as the role of human blood components in bacterial transcriptional phenotypes and the response to vancomycin stress.

## RESULTS

### *S. aureus* forms staphylococcal communities (SCs) in 3D collagen gel matrix assay

To generate staphylococcal communities (SCs) using the *in vitro* 3D collagen gel matrix assay, log-phase *S. aureus* is suspended in a collagen gel solution then seeded into individual wells of a 96 well plate. After gel solidification, overlay media containing key components of human coagulation biology (plasma, fibrinogen, and pro-thrombin) is layered on top of each collagen gel matrix (**Fig 1A**). This 3D suspension, coupled with the presence of human coagulation components, supports SC formation. SCs develop across unique z-planes, which is also observed in SACs in mouse infection models (**Fig 1B**) (22).

**Figure 1.**
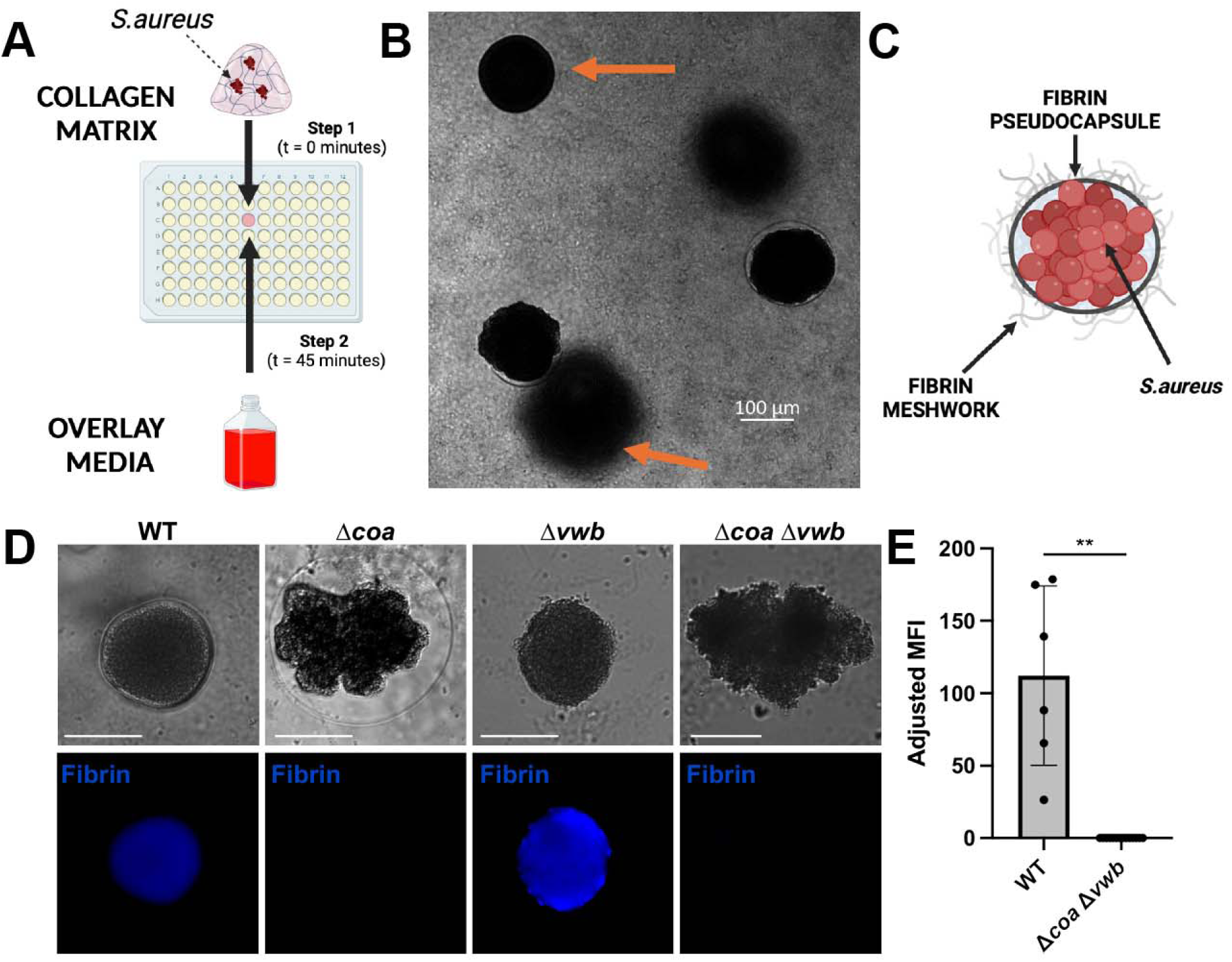
SCs with fibrin barriers form in 3D collagen gel matrix assay in a *coa*-dependent manner. **A)** Schematic of the 3D gel matrix workflow; Created in BioRender. Farhang, P. (2026) https://BioRender.com/8p93erw. Log phase *S. aureus* is suspended in a collagen gel and seeded in a well of a 96-well plate. After solidification for 45 min in the 37°C incubator, overlay media (containing human plasma, human fibrinogen, pro-thrombin, and RPMI+L-glutamine) is added. **B)** Representative image taken of the center of a 3D gel matrix containing SCs at 19h of growth. Arrows are highlighting staphylococcal communities (SCs) growing in different Z-planes. **C)** Schematic of a staphylococcal abscess community (SAC); Created in BioRender. Farhang, P (2026) https://BioRender.com/q8vscr0. **D)** Representative immunofluorescence images of WT, Δ*coa,* Δ*vwb,* and Δ*coa* Δ*vwb* SCs stained for human fibrin. Fibrin was detected using mouse anti-human fibrin antibody and goat anti-mouse AlexaFluor 350 antibody. **E)** Quantification of fibrin staining. Adjusted MFI: total MFI with each strain’s no primary antibody control MFI subtracted (2 biological replicates per strain; 6-14 SCs per strain). Separate batch from 1D. Statistics: Welch’s t-test; ** P<0.01.

Because *in vivo* SACs are defined by their host-derived fibrin structures, we next confirmed the presence of these structures in our SC system **(Fig 1C**). To identify the presence of fibrin structures, WT and mutant (Δ*coa,* Δ*vwb,* or Δ*coa* Δ*vwb*) SCs were grown and probed for the presence of fibrin using immunofluorescence microscopy (38,39). There was a clear presence of fibrin barriers in WT and Δ*vwb* SCs, and an absence in Δ*coa* or Δ*coa* Δ*vwb* SCs (**Fig 1D**). WT SACs have a clear spherical morphology and a more distal structural ring that is vWbp-dependent, and likely represents matrix-associated meshwork (MAM) (**Fig 1D**). Δ*coa* SACs lack a clear spherical morphology yet maintain the distal ring structure, Δ*vwb* SACs lack the distal ring structure but maintain spherical morphology, and Δ*coa* Δ*vwb* SACs lack both features (**Fig 1D, 1E**). Prior literature has observed this distal ring structure and stated that it is comprised of fibrin (although these findings were not supported with immunofluorescence of single deletion mutants), but our fibrin-specific antibody failed to effectively bind it (11,14). This could be due to changes in fibrin structure upon conversion from fibrinogen that mask the antibody epitope when generated by vWbp (as opposed to Coa) in this MRSA strain. Collectively, this data shows that SCs grown in the 3D collagen gel matrix system recapitulate key structural features of SACs observed *in vivo*.

### SCs display temporal regulation and inter-SC heterogeneity in virulence gene expression that can be tracked via fluorescence timelapse microscopy

To determine if we can use this system to track SC gene expression, we incorporated previously described chromosomally-integrated fluorescent reporter strains into the gel matrix system (22). These include *P_agrB_::gfp*, *P_saeP_::gfp*, and GFP^-^ control reporter strains, each in a constitutive mCherry background (*P_sarA_::mCherry*). The GFP^-^ control reporter strain contains *gfp* inserted without a promoter into the same genomic location to control for possible leaky expression and background autofluorescence.

*P_agrB_::gfp* expression was significantly heightened in mature SCs (20h of growth), as expected due to the high density of bacteria within each SC and within a given matrix (**Fig 2A, 2B**). *P_saeP_::gfp* showed expression above baseline but below that of *P_agrB_::gfp*, indicating limited activity in mature SCs (**Fig 2A, 2B**). mCherry levels and SC size were consistent across strains, demonstrating that the fluorescent constructs do not alter SC biology (**Fig 2A, 2C, 2D**).

**Figure 2.**
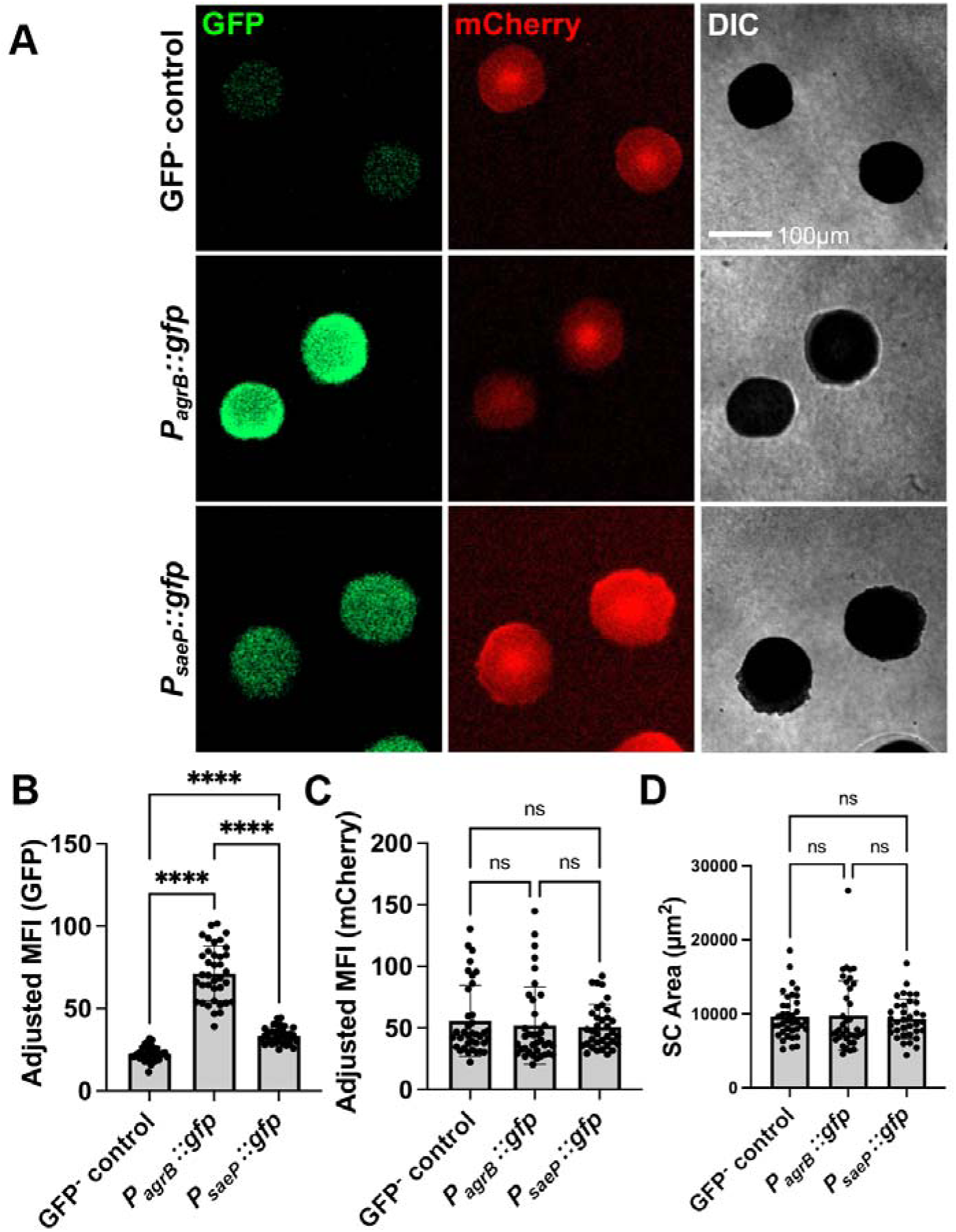
Fluorescent endpoint microscopy detection of *agr* and *sae* activity in mature SCs in the 3D gel matrix. **A)** Representative images of fluorescent reporter strain SCs grown in the 3D gel matrix 20h after overlay media addition. Each strain contains chromosomally-integrated *P_sarA_*::*mCherry* serving as a constitutive signal. **B-C)** Quantification of SC fluorescent reporter signal across a population of SCs grown 20h in the 3D gel matrix. Each dot represents the adjusted mean fluorescent intensity (MFI; Total MFI - background matrix signal) of individual SCs (3 biological replicates per strain; 35-37 SCs per strain). **D)** Quantification of SC area (µm^2^) across a population of SCs grown 20h in the 3D gel matrix. Statistics: **B)** one-way ANOVA and Tukey’s multiple comparisons test; **C & D)** Kruskal Wallace test with Dunn’s multiple comparisons test (at least one of the columns was not normally distributed per Shapiro-Wilk test); **** P<0.0001, ns P>0.05.

In many mouse models of infection, it is not possible to track individual SACs during maturation. SACs within infected mouse tissue are also seeded asynchronously, in the kidney in particular, making it difficult to determine whether SACs represent similar developmental stages at a given endpoint (22). In contrast, our gel matrix system enables the use of automated timelapse microscopy to follow SC development continuously, both across wells and within individual matrices (**Fig 3A**). Target gene expression can also be continuously assessed using fluorescent reporter strains in the matrix system. Using 30 minute automated imaging intervals, we observed that *P_saeP_*::*gfp, P_agrB_*::*gfp,* and GFP^-^ control strains grew exponentially and then gradually plateau after ∼10-15h as SCs fully mature and enter stationary phase (**Fig 3B**). Distinct gene expression patterns were observed based on GFP signal (Adjusted MFI), as *P_saeP_*::*gfp* expression peaked early in SC development and gradually declined as SCs mature, whereas *P_agrB_*::*gfp* expression gradually increased overtime before plateauing in the stationary phase of SC growth (**Fig 3C, 3D**). These trends were confirmed via RT-qPCR (ΔC_T_ values) of bulk RNA isolated from gel matrices (**Fig 3E**). Furthermore, *P_agrB_*::*gfp* fluorescent signal is higher than that of *P_saeP_*::*gfp* in mature SACs (**Fig 3F**), which was corroborated using bulk RNA-seq on matrix-derived RNA at 19.5 hrs (**Fig 3G**). Together, these data demonstrate that microscopy of fluorescent reporter SCs in the gel matrix system provides accurate readouts of gene target transcriptional activity overtime.

**Figure 3.**
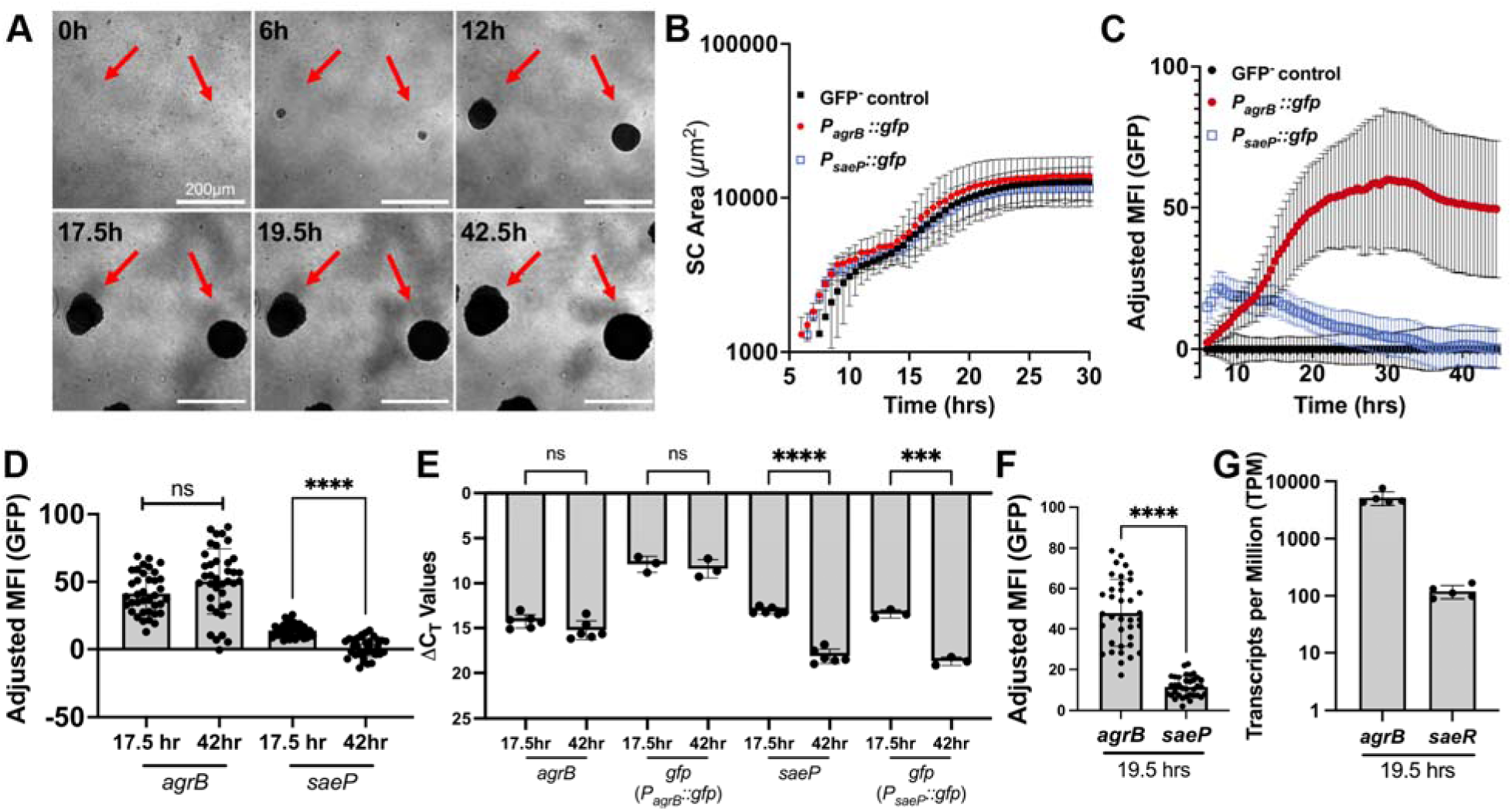
Fluorescent timelapse microscopy shows that *sae* activity decreases while *agr* activity increases in maturing SCs in the gel matrix. **A)** Representative images of SCs grown in 3D gel matrix overtime. Time represents time elapsed after overlay media addition. **B)** Average SC area (µm^2^) per strain per timepoint in 30 min increments (3 biological replicates per strain; 31-38 SCs per strain). **C)** GFP adjusted mean fluorescent intensity (MFI) of fluorescent reporter strain SCs in 30 min increments (3 biological replicates per strain; 31-38 SCs per strain). Average GFP^-^ control GFP signal per timepoint was subtracted from the GFP MFI of the *agrB* and *saeP* reporter strains. **D)** GFP adjusted MFI of individual SCs for each strain at designated timepoints (3 biological replicates per strain; 35-38 SCs per strain). **E)** The centers of individual gel matrices were extracted using a biopsy punch, then processed for RT-qPCR. ΔC_T_ values were generated by subtracting 16S rRNA C_T_ from the target gene C_T_. Each point represents one matrix (6 biological replicates; 3 *P_agrB_*::*gfp,* 3 *P_saeP_*::*gfp*). **F)** GFP adjusted MFI of individual SCs for each strain at 19.5h of growth (3 biological replicates per strain; 35-37 SCs). **G)** Transcripts per million (TPM) from bulk RNA sequencing of whole matrices after 19.5h of growth (5 biological replicates; non-fluorescent WT strain). Statistics: **D-F**) Welch’s t-tests for each pairwise comparison with a Bonferroni multiple comparisons correction to adjust each p value where applicable; **G)** Mann Whitney U test was used due to a non-normal distribution according to a Shapiro-Wilk test; **** P<0.0001, **P<0.01, ns P>0.05.

In addition to measuring a SC population throughout a well, we are able to segregate the size and fluorescent intensity readouts of each individual SC (**Fig 4**). These single-SAC analyses highlight that SACs grown in this system display inter-SAC heterogeneity in size and transcriptional activity, similar to the heterogeneity that has been observed during mouse infection (22).

**Figure 4.**
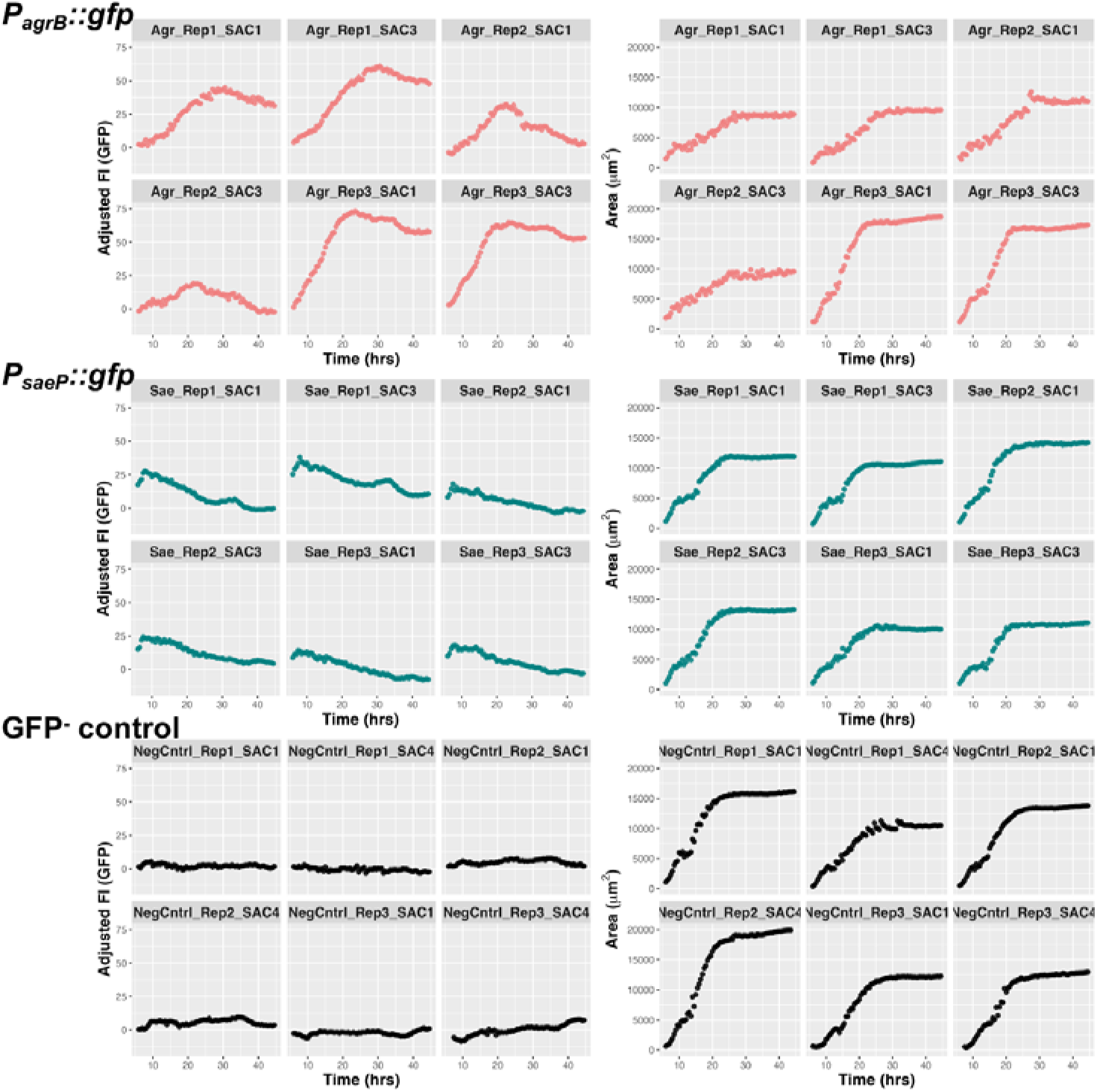
SCs grown in gel matrix display interSC heterogeneity in growth and virulence factor expression. Area (µm^2^) and adjusted GFP fluorescent intensity (FI) of individual SCs from experiments in Fig 3B-C. SCs are identified in the order of reporter strain (i.e. *P_agrB_::gfp* is denoted by “Agr”), biological replicate, and the individual SC number within that biological replicate and strain.

While not a focus of this manuscript, this gel matrix model can also be used to assay host-pathogen interactions, in line with previous iterations of this assay (11,14,36). Primary human neutrophils (PMNs) were co-cultured with nascent SCs and assessed for changes in SC dynamics (**Fig S1A**). While colony forming units (CFUs) were unchanged **(Fig S1B**), SC size was restricted (**Fig S1C**), and *P_saeP_*::*gfp* expression was heightened (**Fig S1D**) in SCs co-cultured with PMNs. These results provide precedent for assaying host-pathogen interactions in this system and offer another avenue for future investigation.

### Fibrin barrier is protective against vancomycin in log-phase SCs

Given *coa* and *vwb* contribute to colony morphology within SCs and fibrin barrier formation (**Fig 1D**), we specifically sought to determine whether the presence of fibrin barriers could impact vancomycin treatment efficacy. Prior mouse infection studies have shown that fibrin barriers protect SAC-residing bacteria, as mutants lacking these structures struggle to establish infection (8,19). Because of the attenuation of these strains (Δ*coa* Δ*vwb*), it has been difficult to assess the specific role of the fibrin barrier during infection and antibiotic treatment. While the exact mechanism of protection has remained elusive, some evidence suggests that the fibrin barriers could restrict drug penetration into fully mature SACs (14).

To test the role of the fibrin barrier in the antibiotic susceptibility of log phase SCs, WT, WT (serum), Δ*coa* Δ*vwb,* and Δ*sae* SCs were grown into late log phase and treated with 50µg/mL vancomycin, the frontline drug used to treat MRSA, for 5h (37,40–42). The Δ*sae* strain was used here as a control since *coa* is expressed downstream of SaeRS activation, while the WT (serum) condition was used as a control meant to phenocopy the Δ*coa* Δ*vwb* strain, since these conditions should not support fibrin barrier formation (11,30). The media used in the WT (serum) condition utilized a similar mixture as the standard gel matrix assay, but lacks fibrinogen and pro-thrombin (see Methods). While all three strains grow similarly in culture, the Δ*sae* strain grows slightly faster than the others in the gel matrix (**Fig S2A-C**). To account for this difference in growth dynamics, the WT, WT (serum), Δ*coa* Δ*vwb* SACs were treated with vancomycin after 8.5h of growth and the Δ*sae* after only 6.5h, ensuring they were treated at analogous late-log phase growth stages.

In log phase planktonic culture, all three strains displayed similar levels of vancomycin susceptibility (**Fig 5A**). In contrast, log-phase WT SCs lacked susceptibility, with no change in CFUs upon vancomycin addition relative to untreated controls, whereas WT (serum), Δ*coa* Δ*vwb* and Δ*sae* mutant SCs saw significant CFU reductions (**Fig 5B, 5C**). Representative DIC images of WT (serum) and Δ*sae,* show similar morphology as Δ*coa* Δ*vwb* SCs, highlighting that the fibrin barriers do not seem to be forming (**Fig 1D, 5D**). To determine if WT SC tolerance was due to the fibrin barriers restricting vancomycin penetration, WT and Δ*coa* Δ*vwb* SCs were stained with the fluorescent BODIPY-vancomycin conjugate after 8.5h of growth, alongside unstained and stained planktonic control cells. Fluorescent microscopy and flow cytometry data indicate that the fibrin barrier restricts vancomycin diffusion into WT SCs, while Δ*coa* Δ*vwb* SCs were uniformly penetrated, similar to planktonic cells **(Fig 5E, 5F, 5G).** These results indicate that the fibrin barrier protects actively growing SCs from vancomycin by serving as a diffusion barrier.

**Figure 5.**
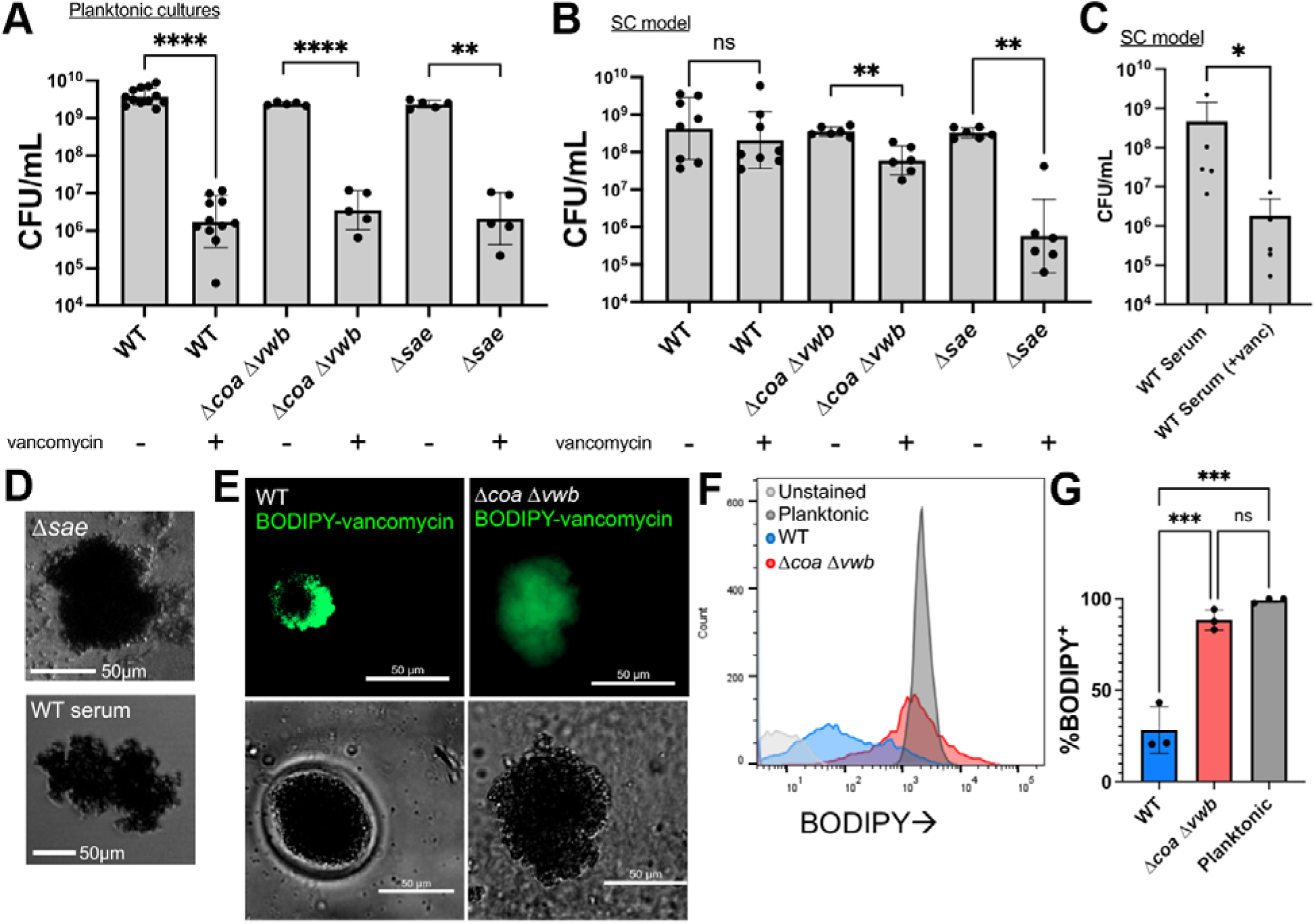
Fibrin barrier protects log-phase SCs from vancomycin. **A)** Vancomycin susceptibility of log-phase planktonic cultures. **B-C)** Vancomycin susceptibility of log-phase SCs. All strains and conditions were treated with 50µg/mL vancomycin for 5h. WT, WT (serum), and Δ*coa* Δ*vwb* SCs were treated at 8.5h of growth, while the faster growing Δ*sae* strain was treated after 6.5h of growth (Fig S2). **D)** Representative images of WT (serum) and Δ*sae* SCs. **E)** Representative images of SCs grown 8.5h in a glass chamber slide to improve imaging resolution. Matrix reagents were scaled appropriately to account for the larger surface area of each well. SCs were stained with 4µg/mL BODIPY-vancomycin. **F)** Representative flow cytometry histograms of each condition after 8.5h of growth stained with BODIPY-vancomycin (unstained control: WT planktonic cells). **G)** Percentage of BODIPY^+^ bacteria for WT and Δ*coa* Δ*vwb* SCs as well as WT planktonic bacteria (3 biological replicates per condition). Statistics: **A-C)** Welch’s t-test or Mann Whitney U test depending on the normality of each column in each pairwise comparison as determined by Shapiro-Wilk test. Multiple comparisons were accounted for using a Bonferroni correction. **G)** one-way ANOVA with Tukey’s multiple comparisons test; ****P<0.0001; *** P<0.001; **P<0.01; ns P>0.05.

### Stationary phase *S. aureus* are transcriptionally responsive to vancomycin when cultured in the presence of human blood components

After observing that fibrin barriers protect log phase SCs from vancomycin penetration, we were interested in understanding if the presence of the fibrin barrier also impacted bacterial transcriptional responses to vancomycin. Log (8.25h) and stationary (14.5h) phase WT SCs were challenged with vancomycin (**Fig 6A, 6B**). To assess the presence of a transcriptional response we used the *P_saeP_*::*gfp* reporter strain, as vancomycin has been shown to activate SaeRS in *S. aureus*, and timelapse microscopy was used to capture transient transcriptional changes (43,44). Despite a clear restriction of log phase SC area in vancomycin-containing matrices, there was no differential expression of *saeP* when compared to untreated conditions (**Fig 6A, 6B**). In contrast, while stationary phase SCs with and without vancomycin displayed no difference in SC size, SCs in vancomycin-containing matrices had an increase in relative *saeP* expression. This was surprising given the conventional wisdom that stationary phase bacteria tend to be less metabolically active and, therefore, less responsive to drugs than their log phase counterparts (45–47).

**Figure 6.**
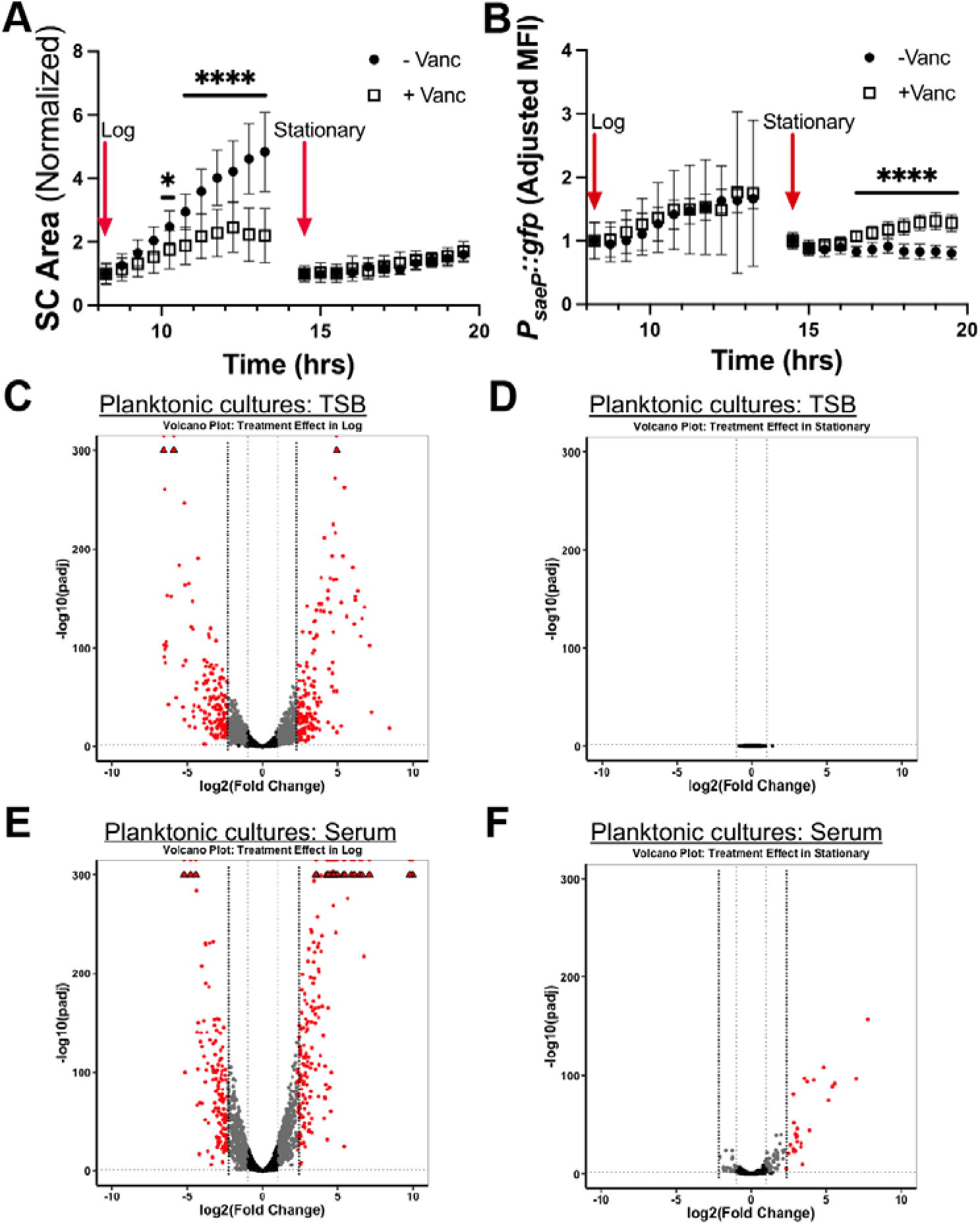
Stationary phase WT SCs respond transcriptionally to vancomycin. A-B) *P_saeP_::gfp* SCs were grown in the gel matrix system for 8.25h (log) or 14.5h (stationary) in triplicate. 0 or 50µg/mL of vancomycin was added in technical replicates of each biological replicate, then imaged overtime at 37°C. SC size (µm^2^) and adjusted mean fluorescent intensity (MFI) were tracked (3 biological replicates; 13-24 SCs per timepoint). **C)** TSB grown log phase planktonic WT treated with 0 or 25µg/mL vancomycin for 2h, fixed, and processed for RNA isolation. **D)** The same experiment as Fig 6C was performed in stationary phase (vancomycin was added after 5.5h of growth). **E)** Serum grown log phase planktonic WT treated with 0 or 25 µg/mL vancomycin for 2h, fixed, and processed for RNA isolation. **F)** The same experiment as Fig 6E was performed with *S. aureus* grown to stationary phase (vancomycin was added after 5.5h of growth). In the volcano plots, black dots represent genes with Padj >0.05 and/or |log_2_(Fold Change)| <1; grey dots indicate genes with Padj < 0.05 and |log_2_(Fold Change)| >1 but < 2.32; red dots indicated genes with Padj < 0.05 and |log_2_(Fold Change)| >2.32. Statistics: **A-B)** multiple unpaired t-tests and multiple comparisons were corrected for using the Holms-Sidak method; **** P<0.0001; * P<0.05.

The presence of a differential transcriptional response in stationary phase SCs was noteworthy, and to determine if this was unique to SCs compared to planktonic growth, we treated WT log and stationary phase TSB grown planktonic *S. aureus* with vancomycin (25µg/mL). This lower concentration (vs. 50µg/mL for SCs) was used to prevent excessive cell lysis, and subsequent controls confirmed that 25µg/mL and 50µg/mL of vancomycin produced similar viability (**Fig S3A**). We then assessed the global transcriptional impact of vancomycin treatment (25µg/mL for 2h) on TSB grown planktonic cells via bulk RNA-seq and observed a robust differential expression of genes in log phase but no significant changes in stationary phase (**Fig 6C, 6D**). Several of the genes with differential transcript abundance upon treatment in log phase align with previously reported findings. For example, *vraR* and *vraX*, part of the *vra* family of genes that are highly responsive to vancomycin and play a role in toxic compound export and the cell wall stress response, both had increased transcript levels upon vancomycin exposure (**Supplemental Table 1)** (44,48). To determine if the different media conditions could be impacting transcriptional responses, we repeated planktonic experiments in human serum (plasma would clot under these conditions). Interestingly, we observed a robust transcriptional response in both log and stationary phase serum-grown planktonic bacteria (**Fig 6E, 6F**), and we confirmed vancomycin susceptibility in both media types (**Fig S3A, S3B).** These data indicate that stationary phase *S. aureus* is transcriptionally responsive to vancomycin when grown in media containing human blood components.

### SCs respond to vancomycin regardless of fibrin barrier presence, but WT and Δ***coa*** Δ***vwb* SCs are transcriptionally distinct**

Through RNA-seq, we determined that stationary phase *S. aureus* respond to vancomycin treatment during culture in human serum components. To determine if the presence or absence of fibrin barriers also impacts the transcriptional response to vancomycin in SCs, WT, WT (serum), and Δ*coa* Δ*vwb* SCs were grown into stationary phase, treated with vancomycin or left untreated, and RNA was isolated for bulk RNA-seq.

Comparison of untreated WT and Δ*coa* Δ*vwb* SCs revealed modest but clear transcriptional differences, as the groups segregated upon PCA analysis (**Fig 7A**). Eight transcripts were increased in relative abundance in WT SCs using a |log_2_(Fold Change)| >2.32 cutoff; notable among these were the increased abundance of V8 protease transcripts (*sspA, sspB, sspC, aur*) (49). These transcripts were also increased in abundance in vancomycin treated WT compared to Δ*coa* Δ*vwb* SCs, alongside *essABC* (type VII secretion system) and several phenol-soluble modulin genes (PSMα 2-4) (**Fig 7B**, **Table 1**) (17,50,51). In addition, Gene Ontology (GO) analysis (with a more inclusive |log_2_(Fold Change)| >1 cutoff) revealed that vancomycin treated Δ*coa* Δ*vwb* SCs are enriched for oxidative stress and protein turnover pathways relative to their WT counterparts. These results indicate, for the first time, that fibrin barriers impact the transcriptome of SCs, and seem to suggest that Δ*coa* Δ*vwb* SC*s* display an increase in stress signatures when responding to vancomycin compared to WT SCs.

**Figure 7.**
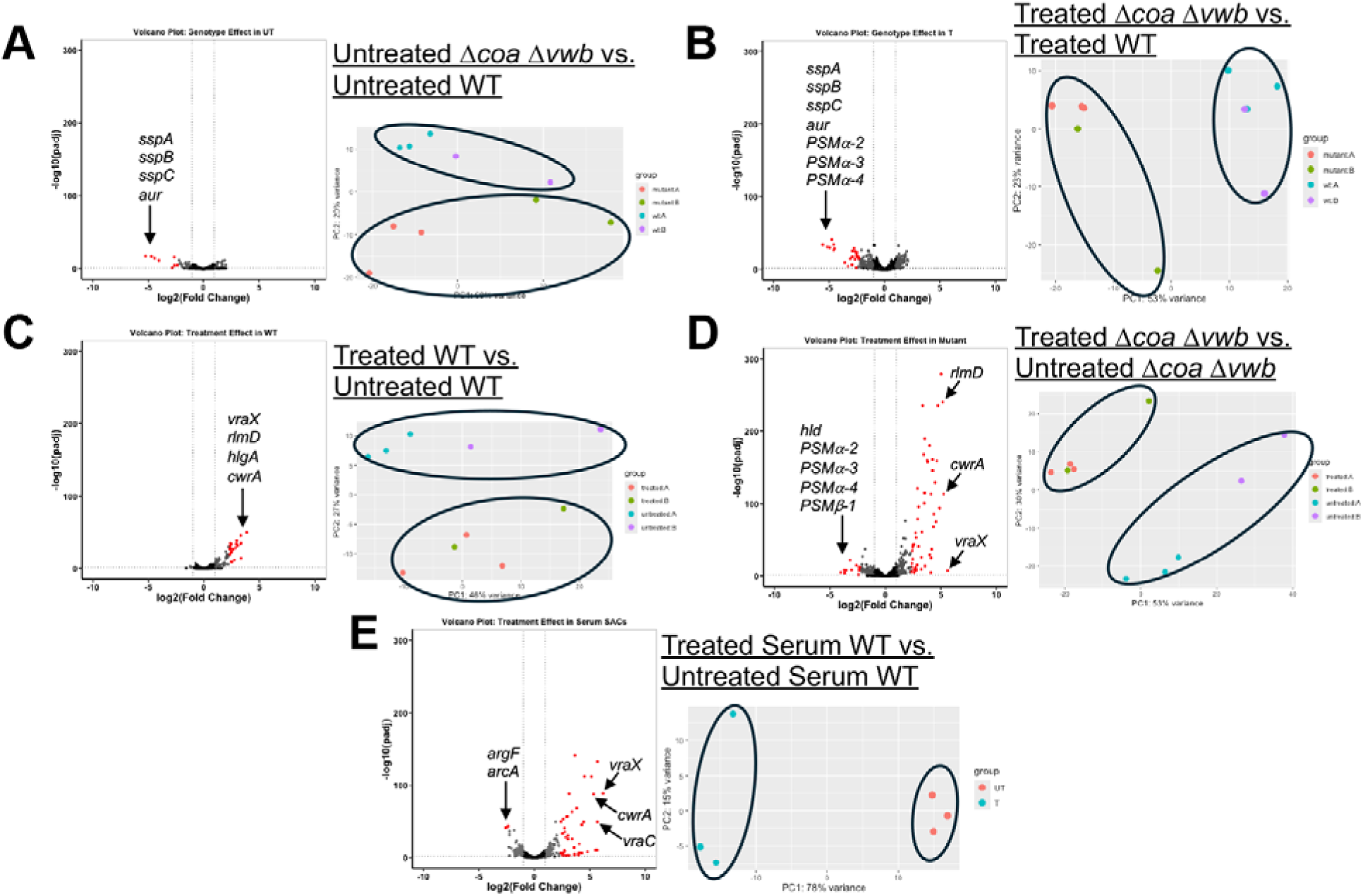
Stationary phase WT SCs respond transcriptionally to vancomycin, whereas. Δ*coa* Δ*vwb* SCs remain in a pre-stressed, unresponsive state. A-E) Differential gene expression analysis of indicated RNA-seq comparisons. Left panels: volcano plots showing genes (baseMean >75) and their -log_10_(Padj) v. log_2_(Fold Change). Black dots represent genes with Padj >0.05 and/or |log_2_(Fold Change)| <1; grey dots indicate genes with Padj < 0.05 and |log_2_(Fold Change)| >1 but < 2.32 (between 2-5 fold); red dots indicated genes with Padj < 0.05 and |log_2_(Fold Change)| >2.32 (≥5 fold). Annotated labels highlight selected genes of interest. Right panels: PCA plots showing separation of comparison groups, with samples colored based on comparison group (ex. treated or untreated) and batch. Ellipses outline comparison groups.

**Table 1.**
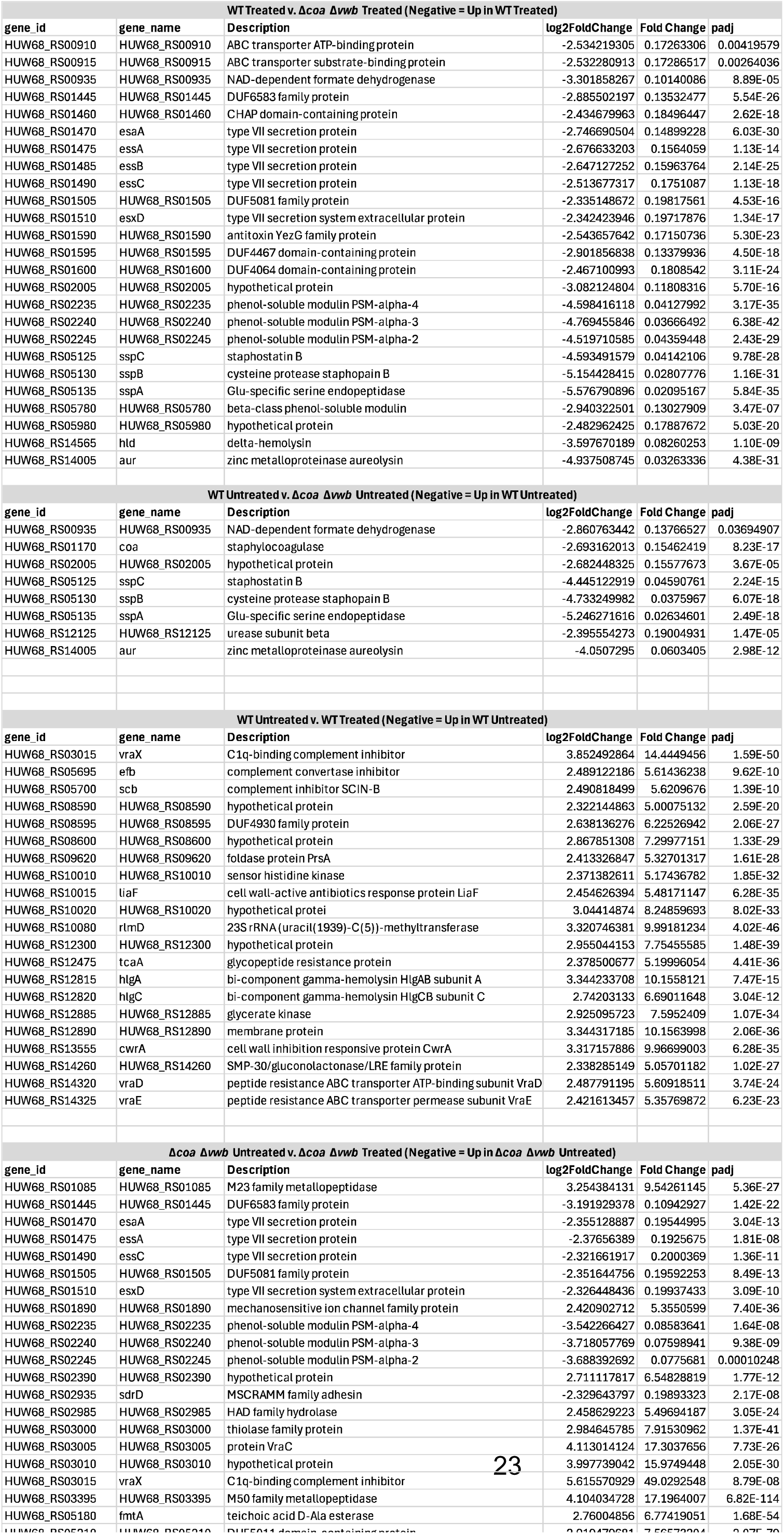
RNA sequencing data of WT, WT (serum), and Δ*coa* Δ*vwb* SCs +/- 50µg/mL Vancomycin (Significant Genes). Comparisons are made between subsets of conditions within the sequencing data set. Each gene is represented by the gene_id listed in the reference genome, the canonical gene_name if it is present, a description of the effector function of the gene product, the log_2_FoldChange, Fold Change, and adjusted p-value for each gene. Cutoffs of log_2_FoldChange +/- 2.32, baseMean >75 reads, and adjusted p-value <0.05 were used as inclusion criteria for genes in each comparison.

When comparing treated to untreated samples within each strain or condition (WT, WT (serum), or Δ*coa* Δ*vwb*), all untreated and treated samples segregated cleanly by PCA, indicating a noticeable response to vancomycin (**Fig 7C, 7D, 7E**). All conditions saw predominantly an increase in transcript levels with treatment, and an increase in the following genes: subsets of *vra* genes (i.e. *vraX, vraR, vraD, vraC*); *hlgA* and *hlgC*, pore-forming toxins (leukocidins); *cwrA*, a cell wall damage responsive gene product that increases cell wall integrity; *rlmD*, a 23S rRNA uridine methyltransferase known to impact antibiotic interaction with ribosomes; and *tcaA*, a glycopeptide resistance protein (17,44,48,52–57). Many of these genes have been closely associated with cell envelope stress and/or vancomycin response in *S. aureus* (*ex. vraR, cwrA,* and *vraX*), while others (ex. *hlgA, hlgC* and *rlmD*) have not been directly tied to these responses (**Fig 7C, 7D, Table 1**). Interestingly, all of these transcripts (*vraX, vraR, tcaA, cwrA, hlgA, hlgC,* and *rlmD* in log phase; *vraX, vraR, tcaA, cwrA,* and *rlmD* in stationary), also increased in vancomycin-treated planktonic bacteria grown in either TSB or serum, underscoring similarities in responses between planktonic and SC-residing bacteria. However, it is important to note there were also many differentially expressed genes unique to each data set (**Supplemental Table 1, Supplemental Table 2)**.

Comparing RNA-seq results by GO analysis, WT, WT (serum) and Δ*coa* Δ*vwb* SCs each show an increase in stress response and phosphorylation (cell signaling) pathways upon vancomycin treatment. These GO terms were also enriched in stationary phase serum-grown planktonic bacteria treated with vancomycin. In WT (serum) and Δ*coa* Δ*vwb* SCs, there was also increased nucleic acid biosynthesis in untreated compared to treated conditions (**Supplemental Table 3**). There were overall more differentially expressed genes in Δ*coa* Δ*vwb* and WT (serum) SCs upon vancomycin treatment (64 and 49 respectively using a |log_2_(Fold Change)| >2.32 cutoff) when compared to WT SCs (20 differentially expressed genes) (**Table 1**), which could be due to drug penetration differences (see Discussion).

Lastly, we utilized the RNA-seq data to compare with fluorescent reporter data in **Fig 6B**, which assessed stationary phase WT *P_saeP_::gfp* SCs treated with vancomycin. Assessing expression of *saeR* and *saeS* in the RNA-seq dataset, both transcripts were significantly increased in vancomycin treated WT SCs relative to untreated, with log_2_ fold changes of 1.17 (*saeR*) and 1.12 (*saeS*) and adjusted p-values of 0.0016 and 0.00026 respectively (**Supplemental Table 2**). These transcriptional changes are consistent with the fluorescent readout from **Fig 6B**, further bolstering confidence in the gel matrix model’s capability of assessing transcriptional spatiotemporal dynamics via timelapse microscopy of fluorescent bacterial reporter strains.

Together, these data indicate stationary phase SCs mount similar transcriptional responses to vancomycin regardless of the presence or absence of fibrin barriers, increasing transcript levels of well characterized cell envelope or vancomycin stress markers as well as several genes not typically associated with these stressors (i.e. *hlgA*). It appears that fibrin barriers do promote a transcriptionally distinct state within SCs. Understanding if there are functional consequences downstream of these strain differences could be important for deepening our understanding of SAC pathogenesis.

### No difference in vancomycin susceptibility, regrowth potential, and PMN cytotoxicity between stationary phase WT and **Δ***coa* **Δ***vwb* SCs

To understand whether the transcriptional differences observed between stationary phase WT and Δ*coa* Δ*vwb* SCs translated into functional differences, we first assessed the impact on vancomycin susceptibility. Stationary phase WT, Δ*coa* Δ*vwb*, and Δ*sae* SCs were treated with 50µg/mL vancomycin for 5h and compared to untreated controls. The Δ*sae* strain was used again as a control since *coa* is expressed downstream of SaeRS activation, and also because a subset of genes in the RNA-seq dataset are expressed downstream of SaeRS (i.e. *hlgA, hlgC*). In contrast with exponential phase results, all three strains lacked vancomycin sensitivity in stationary phase (**Fig 8A**). This was an interesting result, as mature SCs maintain transcriptional responsiveness while exhibiting drug tolerance, and could suggest some of the differentially expressed gene products may have some connection to vancomycin tolerance.

**Figure 8.**
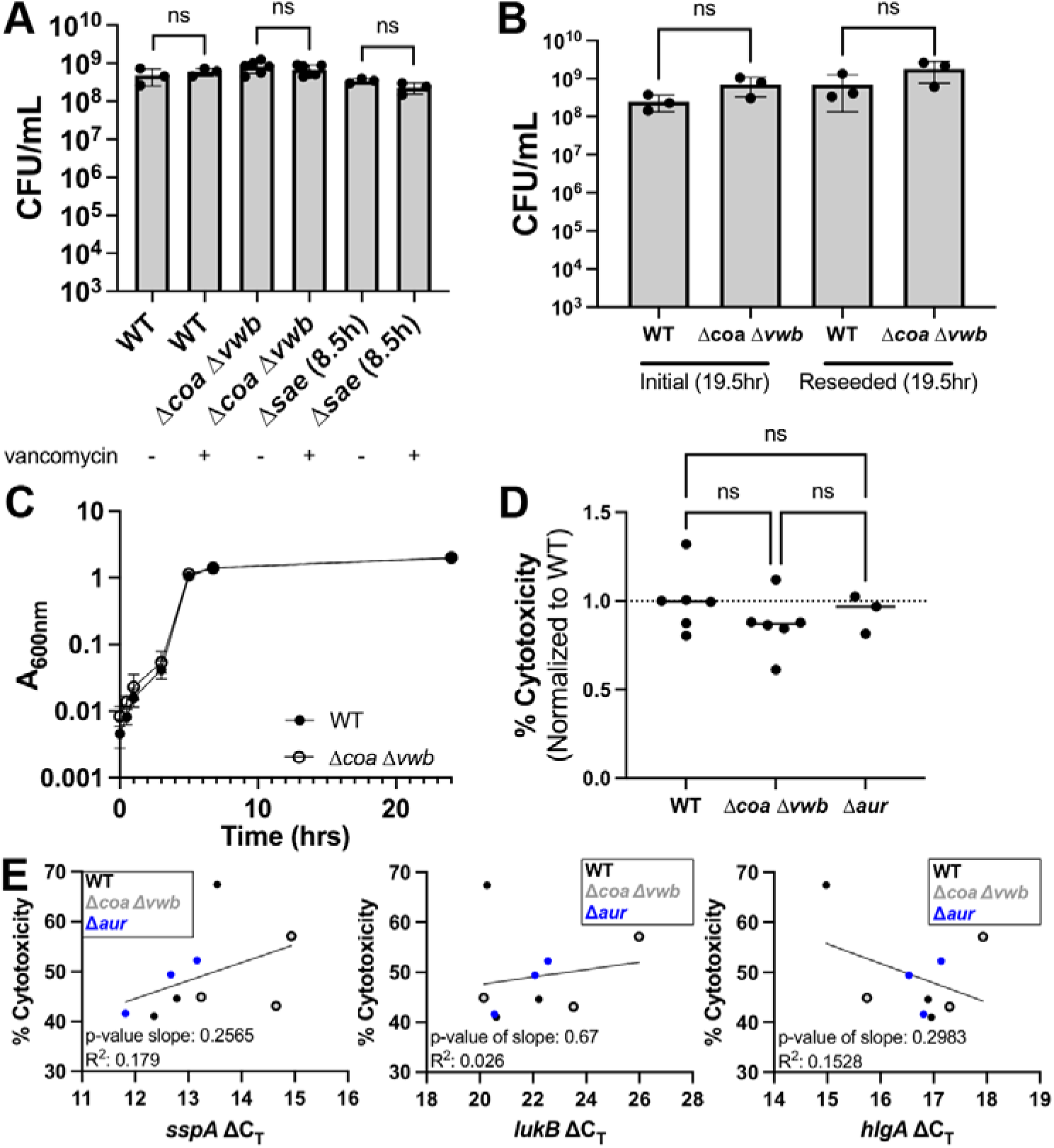
Stationary phase WT and Δ*coa* Δ*vwb* SCs show no differences in vancomycin susceptibility, regrowth potential, or neutrophil cytotoxicity. **A)** Viability of stationary phase WT, Δ*coa* Δ*vwb,* and Δ*sae* SCs treated with 0 or 50 µg/mL vancomycin for 5h. All three strains displayed vancomycin tolerance (3-6 biological replicates per strain). **B)** Reseeding assay. Stationary phase SCs were grown for 19.5h, digested and assessed for CFUs, and reseeded into new matrices. After 19.5h of growth, matrices were again digested and assessed for CFUs (3 biological replicates per strain). **C)** Subset of digested SCs after second 19.5h growth were inoculated into TSB, and A_600_ was tracked overtime (3 biological replicates per timepoint per strain). **D)** Cytotoxicity of PMNs exposed to SC culture supernatants from WT, Δ*coa* Δ*vwb,* and Δ*aur* strains, measured by LDH release. Experiment was repeated across two batches, and % cytotoxicity values were normalized to average value of PMNs cultured with WT SC supernatant within the same batch (3-6 biological replicates per strain). **E)** Simple linear regression comparing toxin transcript levels (ΔC_T_) with corresponding unnormalized PMN % killing (3 biological replicates per strain had both cytotoxicity readout and ΔC_T_ values). Strains are denoted by symbols in panel legend. Statistics: **A)** Welch’s t-test with Bonferroni correction for multiple comparisons; **B)** Welch’s t-tests with Bonferroni correction for multiple comparisons; **D)** one-way ANOVA and Tukey’s multiple comparisons test; ns P > 0.05.

We hypothesized that the stressed phenotype observed in Δ*coa* Δ*vwb* SCs could impact the ability for Δ*coa* Δ*vwb* to resume growth in the context of infection, following dissemination and re-seeding of new infection sites. To test this hypothesis, we grew WT and Δ*coa* Δ*vwb* mutant SCs into stationary phase then reseeded them in new matrices and measured viable counts after 19.5h. There were no differences in CFUs between strains, indicating the Δ*coa* Δ*vwb* transcriptional signature did not come at a fitness cost in its ability to form new mature SCs (**Fig 8B**). In addition to measuring viable counts within SCs, we also inoculated TSB with these WT and Δ*coa* Δ*vwb* reseeded bacteria and assessed growth in liquid culture. We saw no difference in growth between strains via optical density, indicating that stationary phase Δ*coa* Δ*vwb* SCs were able to resume growth with kinetics similar to the WT strain (**Fig 8C**).

We were also interested in determining if transcriptional differences in untreated Δ*coa* Δ*vwb* SCs relative to untreated WT SCs resulted differences in production of components cytotoxic to human neutrophils (PMNs). We probed for transcript levels of toxin expression known to target neutrophils, *lukB* and *hlgA*, as well as the *ssp* operon (17) from SCs and isolated culture supernatant from the matrices. PMNs were isolated from human blood, exposed to culture supernatant from WT or Δ*coa* Δ*vwb* stationary phase SCs, and cytotoxicity was assessed by PMN LDH release. WT samples initially appeared to promote more cytotoxicity than Δ*coa* Δ*vwb*, and an aureolysin mutant (Δ*aur*) was included to determine if V8 protease activity could contribute to any observed effects (58). However, we did not observe significant differences amongst the groups (**Fig 8D**). To assess the expression of the aformentioned genes of interest, RT-qPCR was performed with a subset of biological replicates. Some genes, such as *hlgA* and *lukB* (**Fig 8E**, middle and right panels), did not display a consistent upregulation in Δ*coa* Δ*vwb* SCs across all biological replicates, directionally aligning with the RNA-seq data. Other targets, like *sspA*, showed consistent regulation across all biological replicates, also aligning with the RNA-seq data (**Fig 8E**, left panel: *sspA*). Finally, to understand if there was a correlation between levels of specific transcripts and cytotoxicity, we performed linear regression analyses (**Fig 8E**). Here, lower ΔC_T_ values indicate higher transcript levels, and if heightened transcript levels are linked to increased cytotoxicity, you would expect a negative correlation. We observed this trend with *hlgA*, but this correlation was not significant (**Fig 8E**, right panel).

Finally, to determine if there was an impact of the fibrin barrier on vancomycin sensitivity of *in vivo* SACs, we completed experiments in a mouse model of systemic infection with GFP^+^ WT and GFP^+^ Δ*coa* Δ*vwb* strains (see Methods). Mice were infected retro-orbitally, and at day 4 post-inoculation (p.i.) liver and kidneys were harvested to quantify bacterial load and visualize bacteria by fluorescence microscopy. Day 4 was chosen due to frequency of fully formed SACs at this timepoint. Additional groups of mice were treated with vancomycin (40mg/kg) at day 4 p.i. and tissues were harvested 24 hours later. Surprisingly, we did not observe any difference in bacterial load between the GFP^+^ WT and GFP^+^ Δ*coa* Δ*vwb* strains in either the liver or kidney, or any change in bacterial load following treatment (**Fig 9A, 9B**). To determine if there were differences in *in vivo* SAC morphology, we focused on the % of bacteria that had formed late stage SACs in the kidney at each timepoint, which are stages that have formed a fibrin barrier (**Fig 9C, 9D, 9E**). Again we did not detect differences between strains or differences after treatment, although the % of late stage SACs did seem to increase with the WT strain in the presence of vancomycin, suggesting very little antibiotic sensitivity. These results are consistent with the vancomycin tolerance observed with stationary phase SCs, and may indicate earlier timepoints could reveal increased vancomycin sensitivity.

**Figure 9.**
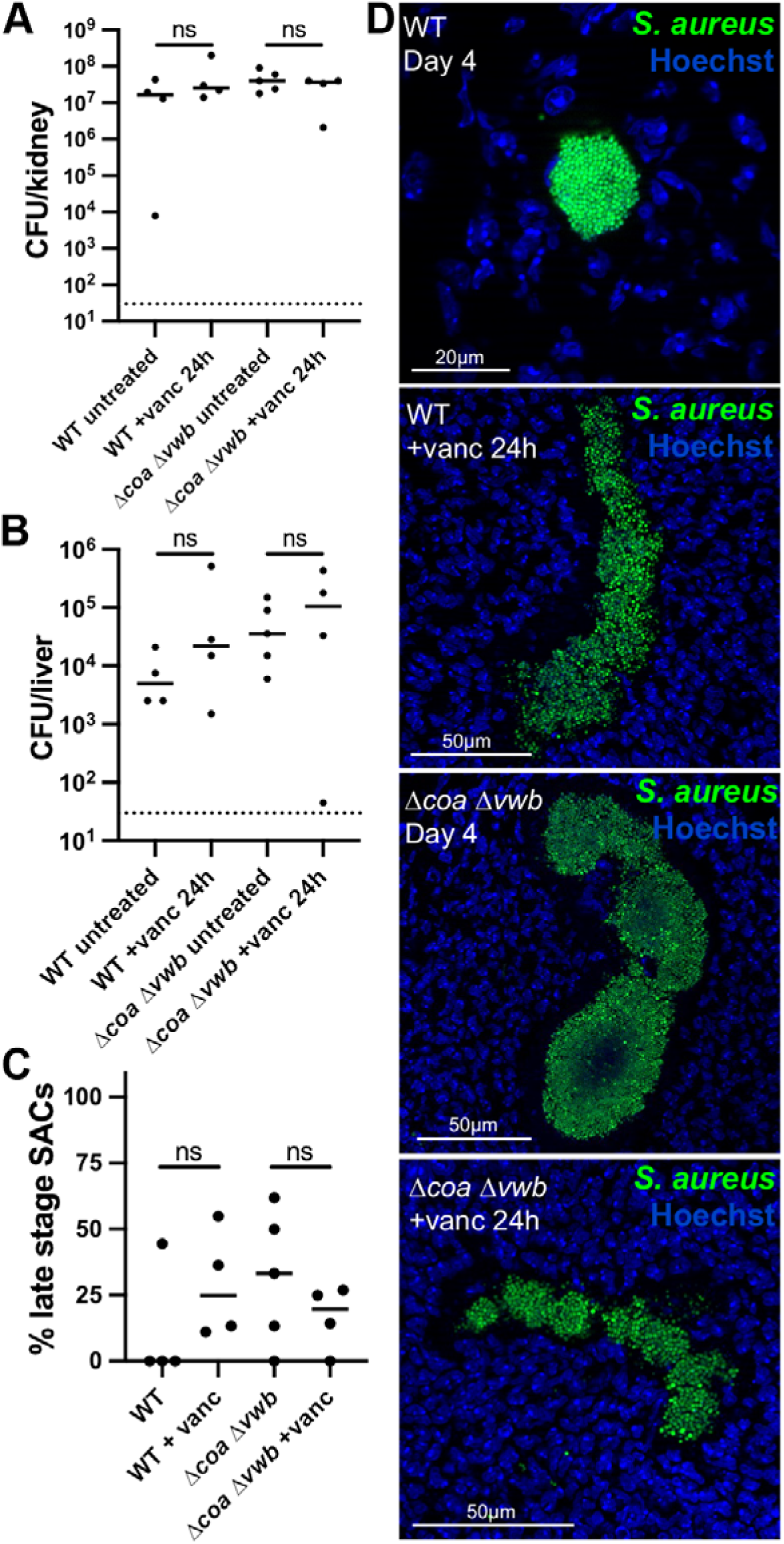
Limited vancomycin susceptibility observed with *in vivo* SACs in the mouse kidney. C57BL/6 mice were inoculated retro-orbitally with 10^6^ CFU of GFP^+^ WT or GFP^+^ Δ*coa* Δ*vwb* strains. Untreated liver and kidney tissues were harvested at day 4 post-inoculation. Subsets of mice were treated with 40mg/kg vancomycin at day 4 and tissues were harvested 24 hours later to quantify bacterial load and visualize bacteria by fluorescence microscopy. **A)** CFU/kidney. Dots: individual mice. Dotted line: limit of detection. **B)** CFU/liver. Dots: individual mice. Dotted line: limit of detection. **C)** % of late stage SACs amongst all bacterial centers present within a single cross-section of kidney tissue. Number of total bacterial centers per tissue ranged between 5 to 31, kidney tissue from all mice represented in panel A is shown here. **D)** Representative images of late stage SACs from each treatment group. Scale bars: either 20µm or 50µm as noted above. Statistics: **A-C)** Mann-Whitney, ns: P>0.05.

Together, while there is compelling evidence showing the fibrin barrier shield actively growing SACs from vancomycin killing and lead to differences in transcriptional responses to vancomycin, these differences did not correspond with measurable changes in vancomycin susceptibility in stationary phase, regrowth potential, or PMN cytotoxicity. Therefore, it appears that the fibrin barrier largely serves as a physical diffusion barrier.

## DISCUSSION

Here we expanded upon prior work to develop a refined 3D collagen gel matrix model for studying staphylococcal community (SC) biology. This iteration offers several novel advantages: i) it utilizes a 96-well plate for high-throughput analyses, ii) supports time-lapse microscopy, and iii) enables recovery of total RNA for bulk transcriptomic analyses. Using this system, we observe that *agr* is expressed as SACs mature, whereas *sae* is expressed early and declines over time. *agr* expression likely increases due to an accumulation of AIP in the local environment, while early Sae activity may be necessary to drive *coa* expression for proper fibrin pseudocapsule formation as SACs mature. To our knowledge, this represents the first high temporal resolution monitoring of these key virulence factors over the course of SC development.

We were interested in utilizing this system to understand fibrin barrier function, and its possible role during antibiotic treatment, with a focus on vancomycin due to its clinical relevance. Prior investigation with a similar model assessed SC-gentamicin interaction, and provided representative images that indicated fibrin barriers restricted diffusion of fluorescently-labeled gentamicin (14). Moreover, mature, stationary phase SCs displayed no susceptibility to gentamicin, unlike isogenic planktonic *S. aureus* treated in parallel. Plasmin pretreatment was used to implicate fibrin barriers, however two limitations complicated interpretation: i) SCs were in stationary phase and therefore less likely to display sensitivity to gentamicin, and ii) plasmin would likely only degrade the outer edges of the fibrin structure and may not completely ablate barrier function (14). To address these limitations, we used *S. aureus* strains with clean deletions of *coa* and *vwb* and treated SCs with vancomycin during active growth. In line with this previous study, we also observed restriction of fluorescently-labeled antibiotic to the periphery of WT SCs. However, as we probed SC antibiotic sensitivity at different growth phases, we found the fibrin barrier protected SCs from antibiotic-mediated killing throughout SC maturation. In contrast, fibrin-deficient mutant SCs saw a complete diffusion of vancomycin and a significant drop in CFUs in log phase compared to untreated controls. These findings together provide compelling evidence that fibrin barriers protect SC-resident bacteria from antibiotics by restricting drug diffusion.

To further probe the role of the fibrin barrier, we compared the transcriptional responses of SCs to vancomycin. Surprisingly, stationary phase SCs were transcriptionally responsive, in contrast to stationary phase planktonic culture grown in TSB, which showed no response to vancomycin (**Fig 6B, 6D**). This led us to determine whether the fibrin barrier or the matrix media (human blood components) was responsible for the stationary phase SC responsiveness. To answer this, we performed several RNA-seq experiments, assessing the vancomycin response in WT *S. aureus* grown in human serum components as well as vancomycin response in stationary phase SCs +/- the fibrin barrier. To our knowledge, this is the first global transcriptional profile (RNA-seq) of SCs, one of the first transcriptional profile of *S. aureus* in serum, and the first transcriptional profile of *S. aureus* in serum +/- vancomycin (59).

We observed that unlike in TSB, planktonic WT *S. aureus* grown in human serum had a transcriptional response to vancomycin in both log and stationary phase. However, the log phase response consisted of many more differentially regulated genes. This indicated that the responsiveness of stationary phase SCs likely stemmed from culturing in human blood components instead of the impact of growth within a community structure.

Although human blood components clearly played a role in the stationary phase phenotype we observed, we also found the presence and absence of the fibrin barrier contributed to the transcriptional response to vancomycin. While there were distinct responses of WT, WT (serum) and Δ*coa* Δ*vwb* SCs to vancomycin, we also noted similarities in terms of cell envelope perturbation-responsive genes. Interestingly, the fibrin deficient samples yielded noticeably more differentially expressed genes. This is likely due to vancomycin penetration differences, and may reflect that more bacteria are exposed to vancomycin.

Despite some similarities, there were noticeable differences in the transcriptome of WT and Δ*coa* Δ*vwb* SCs. When comparing untreated WT and Δ*coa* Δ*vwb* SCs, WT SCs were enriched for the *sspABC* operon and segregated cleanly from mutant SCs upon PCA analysis **(Fig 7A)**. Similar results were obtained when comparing treated WT and Δ*coa* Δ*vwb* SCs, although WT SCs had an increased abundance of transcripts associated with PSMs and a type VII secretion system. Given these SCs were grown in the absence of infection bottlenecks and other stressors associated with a heterogenous host environment, these results could be cleanly attributed to the absence of the fibrin barrier. It would be interesting in future studies to perform single cell RNA-seq or cell sorting with WT SC samples to assess heterogeneity in the transcriptional response. These likely have a population of responding and non-responding SC-residing bacteria due to the physical diffusion barrier presented by the fibrin barrier.

The functional consequences of these differences between WT and fibrin-deficient SACs observed via RNA-seq were not immediately clear. We tested antibiotic susceptibly, reseeding potential, and neutrophil cytotoxicity and observed no difference between SAC strains. Proteomic analysis could help clarify if transcriptional differences are manifesting at the protein level. Heterogeneity in responses may also impact these cytotoxicity results, as the SAC transcriptional changes were not as pronounced in this subset of experiments. Single-cell or small population RNA-seq could also capture intraSAC heterogeneity more fully. Further experimentation of WT and fibrin-deficient SACs in the presence and absence of host factors such as neutrophil co-culture or altered nutritional conditions could better mimic specific tissue niches and may aid in developing translational applications. Together, these would deepen our understanding of how fibrin barriers contribute to SAC development and pathogenesis.

## MATERIALS & METHODS

### Bacterial Strains & Growth Conditions

The *S. aureus* WT USA300 LAC isolate was used as the parent background for all experiments in this study, including reporter and deletion strains. These strains have been previously described (22,38,39,42,58). For mouse infections, GFP-expressing derivatives of LAC WT and Δ*coa* Δ*vwb* mutant strains were used. The GFP+ LAC WT strain has been previously described (60). The GFP+ Δ*coa* Δ*vwb* strain was generated by transducing the GFP plasmid from the GFP+ LAC WT strain into the Δ*coa* Δ*vwb* mutant using bacteriophage phi11, as described previously (61). Transductants were selected on media supplemented with chloramphenicol (10µg/mL). *S. aureus* was grown on tryptic soy agar (TSA) plates at 37°C to isolate single colonies. Overnight cultures were inoculated with a single bacterial colony into tryptic soy broth (TSB) and incubated at 37°C with rotation for 13-20 hrs. Back-dilutions were made in fresh TSB and were placed again on the 37°C rotor for the indicated time. Fluorescent reporter strains contain a kanamycin resistance cassette and were cultured with 50µg/mL kanamycin.

### 3D Collagen Gel Matrix Assay

Overnight cultures of *S. aureus* were back-diluted 1:100 into fresh TSB and grown 2h to exponential phase. The culture was then further diluted 1:10,000 and 50µL (∼2,500 cells) was pelleted at 20,000 x g. The bacterial pellet was resuspended in a 20µl solution of rat tail collagen (ThermoFisher Scientific, cat no. 1048301) and RPMI + L-glutamine (ThermoFisher Scientific, cat no. 11875093) with a final concentration of 3.7mg/mL collagen. The suspension was pipetted into the center of a well in a 96-well plate (Greiner, cat no. 655180) and placed at 37°C for 45 min to allow the collagen matrix to solidify. 48µL overlay media from a master mix consisting of 25µL 10% pooled human plasma reconstituted in PBS (phosphate-buffered saline), 15µL of 10mg/mL human fibrinogen (Millipore Corp, cat no. 341576) reconstituted in PBS, 4µL of 53µg/mL prothrombin (Invitrogen, cat no. RP-43087) reconstituted in 50% glycerol, and 8µL of RPMI + L-glutamine was added onto each matrix. The plate was placed at 37°C until usage in downstream assays. We used a humidified environment for all incubation steps. In this manuscript, the time of growth within the matrix is listed as the time post overlay media addition.

### Plasma Sources

Two different plasma sources were used in experiments due to plasma sourcing/supply chain issues. All relevant experiments except the flow cytometry, fibrin immunofluorescence representative images, quantitative portions of neutrophil co-culture experiments, and second batch of cytotoxicity assay were conducted using lyophilized pooled human citrated plasma from Millipore-Sigma (cat no. P9523) reconstituted in PBS (100mg powder per 1mL PBS). The aforementioned experiments were conducted using pooled human citrated plasma from Innovative Research Inc (cat no. IPLAWBNAC50ML). This plasma source was delivered frozen at –80°C. For flow cytometry, quantitative portions of neutrophil co-culture experiments, and second batch of cytotoxicity assay, this plasma was lyophilized for 48h, then reconstituted in PBS (100mg powder per 1mL PBS) and filtered using a 5µm filter. For fibrin immunofluorescence representative images, plasma was freeze thawed from single use aliquots and filtered using a 5µm filter. SACs grown under various plasma preparations were assessed via microscopy and RT-qPCR, ensuring similar morphology, growth kinetics, and transcriptional profile.

### Serum Source

Experiments were conducted using pooled human serum ‘off the clot’ from Innovative Research Inc (cat no. ISER50ML), freeze thawed from single use aliquots. Data from Supplemental 3A used serum that was filtered using a 5µm filter (required a larger volume), all other experiments were filtered using a 0.45µm filter. Filtration had no impact on endpoint CFU/mL. Experiments involving serum utilized the same media mixture as the gel matrix with the following exceptions: serum instead of plasma, PBS instead of fibrinogen dissolved in PBS, and 50% glycerol instead of pro-thrombin dissolved in 50% glycerol. The reference to “serum” in the body of the manuscript refers to this overlay media mixture.

### Microscopy

The 96-well plate containing matrices with overlay media was placed in the Zeiss Axio Observer 7 inverted fluorescent microscope. Images were captured with an Axiocam 702 mono camera (Zeiss) and processed using ZEN3.10 software. Using a 5X objective, the field of view was adjusted to the center of a well and then further adjusted along the z-plane to the midline of the matrix. For timelapse microscopy, the chamber was pre-warmed to 37°C, and image acquisition was automated to take images across all matrices of a given 96-well plate every 30 minutes.

### Microscopy Quantification

Only SACs in focus, fully in the plane of view of a given image, and not touching other SACs, were included in quantification. Fluorescent signal was quantified using Volocity software (Quorum Technologies Inc). Except for the fibrin immunofluorescence experiment, background signal from SAC-less regions of a given matrix were subtracted from all fluorescence quantification. For experiments where individual SACs were tracked overtime, background signal from the area immediately adjacent to SAC was subtracted from each SAC; for experiments where individual SACs were not tracked overtime (i.e. the population was tracked overtime) the background signal was taken from one or two SAC-less areas that were representative of overall background signal in the matrix.

### Fibrin Immunofluorescence

Gel matrices were grown containing either WT or Δ*coa*Δ*vwb S. aureus*. Wells were fixed with 4% PFA for 30min at room temperature after 12.5h (Fig 1D) or 18h (Fig 1E) of growth. Matrices were washed with PBS then blocked with 2% BSA solution for 1hr at room temperature. Matrices were washed with PBS then incubated overnight at 4°C with 100µL of a 1:100 dilution of anti-fibrin primary antibody (Sigma-Aldrich, cat no. MABS2155) in 2% BSA. Matrices were washed with PBS then incubated at room temperature for 1hr with 100µL of a 1:500 dilution of secondary antibody (ThermoFisher Scientific, cat no. A21120) in 2% BSA. Matrices were washed with PBS.

### RNA Isolation

Samples fixed in 4% PFA were processed for bulk RNA isolation. Fixation lasted for 30 min at 4°C for experiments validating matrix fluorescent protein expression, remaining experiments were either fixed for 30min at room temperature or 4°C overnight. For experiments validating matrix fluorescent protein expression, a 2mm biopsy pen was used to extract central contents of the gel matrices and placed in a 1.5mL microcentrifuge tube. Matrices were washed with PBS and centrifuged to isolate matrix contents, then resuspended in a PBS solution containing 1U collagenase enzyme. Matrices were incubated until visibly dissolved (enough to suspend into solution in subsequent steps), typically 30min at 37°C. Matrices were washed with PBS and centrifuged, then resuspended in 1mg/mL lysostaphin in 100mM Tris HCl (Millipore Sigma, cat no. L7386) and incubated at 37°C 30min. RNA isolation was performed using reagents from the TRIzol Max Bacterial RNA Isolation Kit (Invitrogen, cat no. 16096020) on the digested samples, followed by the Turbo DNA-free kit (Invitrogen, cat no. AM2238) to reduce gDNA contamination, according to manufacturer’s protocols.

For all other experiments, RNA from 4% PFA fixed samples was isolated using the RNeasy FFPE Kit (QIAGEN, cat no. 73504) according to manufacturer’s protocols. For gel matrices, mechanical disruption with sterile rotors and collagenase digestion were used to digest samples. This matrix digestion (or fixed planktonic samples) was then spun and resuspended and incubated in 1 mg/mL lysostaphin in 100mM Tris HCl no more than 30min at 37°C. RNaseOUT (Invitrogen, cat no. 10777019) was used during this digestion step in early iterations of this method. The RNeasy FFPE kit was then used to isolate RNA from the resulting lysate. The On-column DNase digestion kit (QIAGEN) was used according to manufacturer’s protocols. If residual gDNA was detected by RT-qPCR controls, the Turbo DNA-free kit was subsequently used according to manufacturer’s protocols.

### cDNA Synthesis, and RT-qPCR

To generate cDNA, the ProtoScript II First Strand cDNA Synthesis Kit was used (New England Biolabs, cat no. E6560S). Resulting cDNA was diluted in 80µL nuclease-free water and stored at -20°C. For the qPCR reactions, primers for *agrB*, *saeP*, *vraR, hlgA, sspA, lukB,* and 16s rRNA were used in concert with the PowerUp SYBR Green Master Mix (Applied Biosystems). Signal from the reaction was captured on a StepOnePlus System (Thermo Fischer Scientific). Relative quantification was assessed using ΔC_T_ values (16s C_T_ value subtracted from target C_T_ values of corresponding sample). All kits and reagents were used according to the manufacturers’ protocols.

### Antibiotic Susceptibility

To assess antibiotic susceptibility in planktonic culture, overnight cultures of *S. aureus* strains were back-diluted 1:100 in fresh TSB and incubated 2h at 37°C with rotation for log phase samples. Cultures were then diluted 1:10 in TSB in technical replicates (∼5 x 10^7^ CFU/mL). Technical replicates were given 50, 25 or 0 µg/mL vancomycin. Samples were incubated at 37°C with rotation for 5h then plated for viable counts. To assess antibiotic susceptibility in matrices, matrices were prepared and grown in technical replicate pairs for the indicated amounts of time. One matrix in each technical replicate pairing was treated with 50µg/mL vancomycin while the other was left untreated. Treatment lasted 5h, then matrices were digested with collagenase until the matrix was visibly dissolved, approximately 30 minutes at 37°C. Matrices were then plated for viable counts.

### BODIPY-Vancomycin Staining & Flow Cytometry

Matrices were grown for 8.5h, washed 1x with PBS, then incubated +/- 4µg/mL BODIPY-vancomycin conjugate (ThermoFisher Scientific, cat no. V34850). Matrices were incubated for 10 minutes at 37°C, washed, then fixed with 4% PFA. Matrices were mechanically disrupted with sterile rotors and digested with collagenase for approximately 30 minutes at 37°C. Digested matrices were then placed in FACS tubes and run through the BD FACSymphony A3 flow cytometer.

### RNA Sequencing Experiments

For planktonic experiments, WT *S. aureus* overnight cultures were back-diluted 1:100 in TSB or serum and grown for 2h (log) or 5.5h (stationary) and placed at 37°C (3 biological replicates per growth phase; for TSB experiments growth phases were conducted in different batches; for serum experiments all conditions were done in a single batch). Technical replicates per biological replicate were then treated with 0 or 25 µg/mL vancomycin for 2h at 37°C, fixed, and processed for RNA isolation. TSB experiments were done in glass tubes and placed on a rotor during incubations to mimic standard laboratory cell culture conditions. Serum experiments were done in a 96 well plate and not placed on a rotor to mimic SAC experiments and minimize volume of filtered serum needed. Bacteria in serum were grown in glass tubes to see if there were notable differences in growth with the 96 well plate format, and no major differences in growth dynamics were observed by A_600._

For SAC experiments, WT, WT (serum) and Δ*coa* Δ*vwb* SACs were grown to stationary phase in the gel matrix model and treated with 0 or 50 µg/mL vancomycin for 5h prior to RNA isolation. WT and Δ*coa*Δ*vwb* conditions were grown for 14.5h prior to treatment across 2 batches (3 biological replicates per condition in Batch A; 2 biological replicates per conditions in Batch B). WT (serum) conditions were grown for 25.5h prior to treatment in 1 batch.

### RNA Sequencing and Analysis

For RNA-seq, Genewiz (from Azenta Life Sciences) performed library preparation and paired end Illumina sequencing on total RNA samples. Downstream sequencing analysis and differential gene expression analysis was performed by the Davis Lab. The following software packages were used for each step: quality control was assessed using FASTQC, adaptor trimming using the bbduk, alignment was done using bowtie2, hit counts were tallied using featureCounts, and differential gene expression analysis was done via DESeq2 (62). The following reference genome was used for the analysis: ASM1547557v1.

### Reseeding Assay

Matrices were grown for 19.5h, then digested with collagenase for 15 minutes at 37°C. The digestion was pelleted and resuspended in 1mL TSB, then diluted 1:1000, and 50µL of the dilution was spun down and incorporated into new gel matrix (∼2,500 CFUs). Fresh overlay media was added immediately upon matrix seeding. Matrices were grown another 19.5h then digested with collagenase again. Aliquots were plated for viable counts, while the rest was diluted 1:10 in fresh TSB and incubated at 37°C with rotation. Absorbance (600nm) measurements were taken at the indicated timepoints from these planktonic cultures.

### Agarose Pads

Overnight cultures of *P_saeP_*::*gfp S. aureus* were split into technical replicate pairs and given 50µg/mL vancomycin or left untreated and placed at 37°C with rotation. 20µL of solution was sampled and fixed in 4% PFA per tube 0, 2, and 5h after antibiotic addition. Agarose pads were prepared to immobilize bacteria and assess bacterial fluorescence by microscopy. 25µL of 1% agarose dissolved in PBS was pipetted onto a standard microscope slide, covered by a coverslip, and left to solidify at room temperature for 30 minutes. Once the agarose solidified, the coverslip was removed and ∼8µL of fixed bacteria were added. The pipetted bacteria were then covered by a fresh coverslip and imaged at 63X magnification using the fluorescent microscopy setup described above (63,64).

### PMN Isolation & Gel Matrix Incorporation

Buffy coats were obtained from the American Red Cross (ARCBS IRB: 2020-017). Samples were spun at 800 x g for 10min to separate red blood cells (RBCs). The immune cell-containing supernatant was then placed on a Mono-Poly Resolving Medium (MP Biomedicals, cat no. 1698049) in a 1:1 ratio and centrifuged for 45min at 800 x g. The granulocyte layer was then removed and processed to remove remaining RBCs. 1mL Red Blood Cell Lysis Buffer (Roche, cat no. 77443200) was mixed with 500µL granulocytes for 10 minutes at room temperature. Contents of this mixture were spun at 500 x g for 5 min, and the resulting pellet was suspended in another 1mL of Red Blood Cell Lysis Buffer. After another 500 x g for 3 minute spin, purified neutrophils were resuspended in media and incorporated into the appropriate gel matrices (∼100,000 neutrophils). Neutrophils were added either 40 minutes (representative images) or 2h (quantitative analysis) after seeding. For each isolation, the purity of immune cell preparations was confirmed by flow cytometry. Neutrophils were identified as CD66b^+^CD49d^-^ using a CD66b monoclonal antibody (G10F5, APC; Thermo Fisher Scientific, cat no. 17-0666-42) and a CD49d monoclonal antibody (9F10, PE; Thermo Fisher Scientific, cat no. 12-0499-42).

### LDH Cytotoxicity Assay

WT, Δ*coa* Δ*vwb*, and Δ*aur* SACs were grown in gel matrices for 19.5h using 150µL of RPMI + L-glut without phenol red as the RPMI source. Matrices were isolated from wells, placed in microcentrifuge tubes, and spun at 20,000xg. The supernatant was removed, placed in separate tubes, and stored at 4°C. Once PMNs were isolated, we seeded relevant wells of a V-bottom plate (Greiner, cat no. 651180) with ∼500,000 PMNs in 140µL matrix supernatant. As a positive control, PMNs were resuspended in 140µL matrix supernatant with Triton X-100. As a negative control, PMNs were resuspended in RPMI + L-glut without phenol red. To account for media background signal, RPMI + L-glut without phenol red and matrix supernatant were placed in wells without PMNs. The V-bottom plate was incubated for 30 minutes at 37°C with 5% CO_2_, then spun at 400xg for 5 min. The top 100 µL of each well was transferred to a flat bottom 96-well plate and 100 µL of LDH reagent (Cell Signaling Technology, cat no. #37291) was mixed in each well, followed by incubation for 30min at 37°C, protected from light, with shaking. A_490nm_ was quantified as a readout for LDH release.

### Murine Infection Model

All animal experiments were approved by the Johns Hopkins University Institutional Animal Care and Use Committee (Protocol#: MO23H310). *S. aureus* frozen stocks in 10% glycerol (prepared after 2h growth from a 1:50 back-dilution of overnight cultures) were washed and diluted in sterile PBS to 10^6^ CFU in 100µL for inocula. 6- to 8-week-old female C57BL/6 mice (Jackson Laboratories) were inoculated intravenously via the retro-orbital (under isoflurane anesthesia) route. At the indicated timepoints post-inoculation, mice were euthanized using a lethal dose of isoflurane, followed by cervical dislocation, and the liver and kidneys were harvested. Left kidneys were homogenized for CFU enumeration, and the right kidneys were fixed in 4% PFA overnight at 4°C for histology. At day 4 post-inoculation, subsets of mice were treated with 40mg/kg vancomycin in 100µl PBS by intraperitoneal injection.

### Fluorescence Microscopy of Mouse Tissue

After fixation, the right kidneys were frozen-embedded in O.C.T. compound (Tissue-Tek, VWR) and stored at −80°C. 10µm sections were cut using a cryostat microtome (Microm HM 505e) and mounted on charged microscope slides (HistoBond, VWR). Sections were thawed in PBS at room temperature (RT) and stained with Hoechst (1:10,000 dilution in PBS) for 15 min. Coverslips were mounted with ProLong Gold (Invitrogen). One section per mouse was imaged using a Zeiss Axio Observer 7 inverted fluorescent microscope with a 63× oil objective and Apotome.2. Images were captured with an Axiocam 702 mono camera (Zeiss) and processed using ZEN3.10 software. SACs were analyzed by stage as previously described (22), and the % of late stage (stages 3 and 4) SACs in each tissue were shown.

## ACKNOWLEDGMENTS

We thank the Davis, Archer, Pekosz, Thompson, and Klein labs and Dr. Erin Goley for constructive feedback and suggestions. Special thanks to Dr. Alexander Horswill and Dr. Jakub Kwiecinski for their expertise on prior iterations of the 3D collagen gel matrix system used in this manuscript. Their feedback and suggestions as the project was being crafted were very helpful. Thank you to the labs of Dr. Victor Torres and Dr. Alexander Horswill for providing bacterial strains used in this study. We also thank the JHSPH Flow Cytometry Core. This work was supported by the National Institutes of Health (AI159473 to KMD) and the Burroughs Wellcome Fund (1180500 to KMD). The funders had no role in study design, data collection and analysis, decision to publish, or preparation of the manuscript. The authors received no specific funding for this work.

## SUPPLEMENTAL DATASETS

**Supplemental Dataset Table:** All data points plotted in the main manuscript and Supplemental Figures. Each tab represents one Figure.

**Supplemental Fig 1. Primary neutrophils impact SC biology in the gel matrix system. A)** Representative images of SCs grown with or without 100,000 primary human neutrophils (PMNs), imaged 20h after overlay media addition. PMNs were added after 40 min of SC growth. **B-D)** *P_saeP_::gfp* SCs were grown in gel matrix system (3 biological replicates). For each biological replicate, one technical replicate received 100,000 PMNs after 2h of SC growth while another used none. Endpoint analysis comparing CFUs per matrix (B), average SC area (µm^2^) (C) and GFP adjusted MFI signal (D) were conducted after 6h of SC growth. Statistics: **B-C)** Welch’s t-test; **D)** Mann Whitney U test (depending on the normality of the groups being compared according to the Shapiro-Wilk test); ** P<0.01, *P<0.05, ns P>0.05.

**Supplemental Fig 2. *S. aureus* strain growth dynamics in planktonic culture vs. in gel matrix system. A)** Blank-subtracted A_600_ measurements of *S. aureus* strains overtime. Overnight cultures were back diluted 1:100 into TSB and grown on a 37°C rotor. **B)** Bacterial load (CFU/mL) was measured at a subset of timepoints from cultures setup in (A). **C)** Gel matrices were grown and visualized using timelapse microscopy. Total bacterial area (µm^2^) in the center of each matrix was assessed per strain per timepoint (3-5 biological replicates per strain).

**Supplemental Fig 3. *S. aureus* planktonic response to different dosages of vancomycin. A-B)** WT overnight cultures were diluted 1:100 in TSB for 2h on 37°C rotor. These cultures were further diluted 1:10 in **A)** TSB or **B)** serum overlay media and treated with 0, 25, or 50µg/mL vancomycin for 5h on a 37°C rotor before taking raw A_600nm_ measurements. Statistics: **A-B)** one-way ANOVA with Sidak’s multiple-comparison correction; ****P<0.0001; ns P>0.05.

**Supplemental Table 1. RNA-seq data of WT Planktonic Bacteria in Log or Stationary Phase +/- 25µg/mL Vancomycin (All Genes).** Comparisons are made between subsets of conditions within the sequencing data set (one comparison per tab). Each gene is represented by the gene_id listed in the reference genome, the canonical gene_name if present, a description of the effector function of the gene product, the log_2_FoldChange, Fold Change, and adjusted p-value. Only genes with baseMean>75 reads are listed. Treatment started after 2h (log) or 5.5h (stationary) of growth and lasted 2h. For TSB data, 3 biological replicates per growth phase across 2 batches (1 per growth phase). For serum data, 3 biological replicates per growth phase across 1 batch.

**Supplemental Table 2. RNA-seq data of WT, WT (serum), and** Δ***coa*** Δ***vwb* Stationary Phase SCs +/- 50µg/mL Vancomycin (All Genes).** Comparisons are made between listed subsets of conditions within the sequencing data set (one comparison per tab). Each gene is represented by the gene_id listed in the reference genome, the canonical gene_name if present, a description of the effector function of the gene product, the log2FoldChange, Fold Change, and adjusted p-value. Only genes with baseMean>75 reads are listed. For plasma conditions, treatment started after 14.5h of growth and lasted 5h (5 biological replicates per condition across 2 batches). For serum conditions, treatment started after 25.5h of growth and lasted 5h (3 biological replicates per condition across 1 batch).

**Supplemental Table 3. Gene Ontology Data for Stationary Phase SC RNA-seq Data.** Gene Ontology (GO) Biological Process (BP) enrichment was performed on genes differentially expressed from RNA-seq data using the topGO package.Enrichment was assessed separately for significantly up and down genes (adjusted p value < 0.05, |log_2_FoldChange| >1, and baseMean >75 reads) using the topGO package with the weight01 algorithm and Fisher’s exact test. The gene universe was restricted to genes detected in the RNA-seq dataset and annotated with at least one GO term. Shown are the top 20 enriched GO terms per analysis; listed genes represent those contributing to each enriched term. Weighted Fischer output is unadjusted.

