## Supplementary figures and images for "Fibrin barriers limit vancomycin penetration into staphylococcal communities and impact *S. aureus* transcriptional responses"

### Supplemental Fig 1

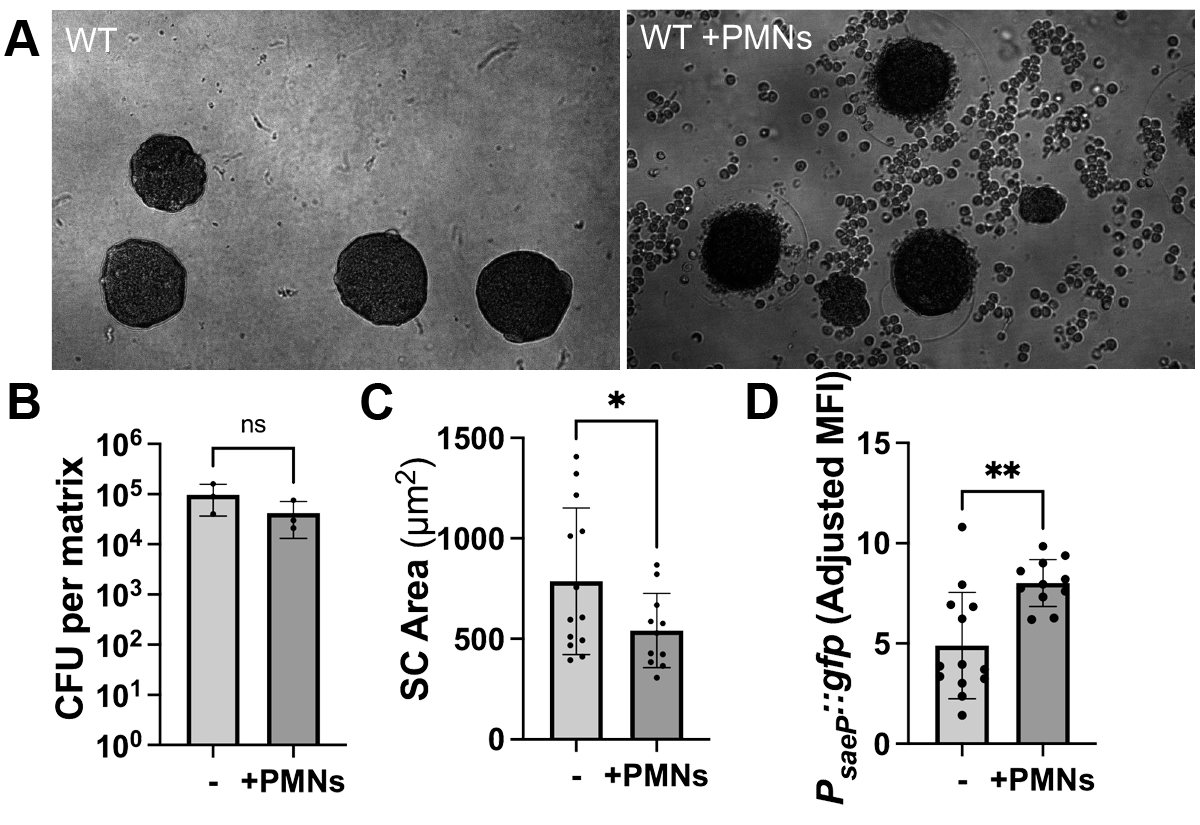

### Supplemental Fig 2

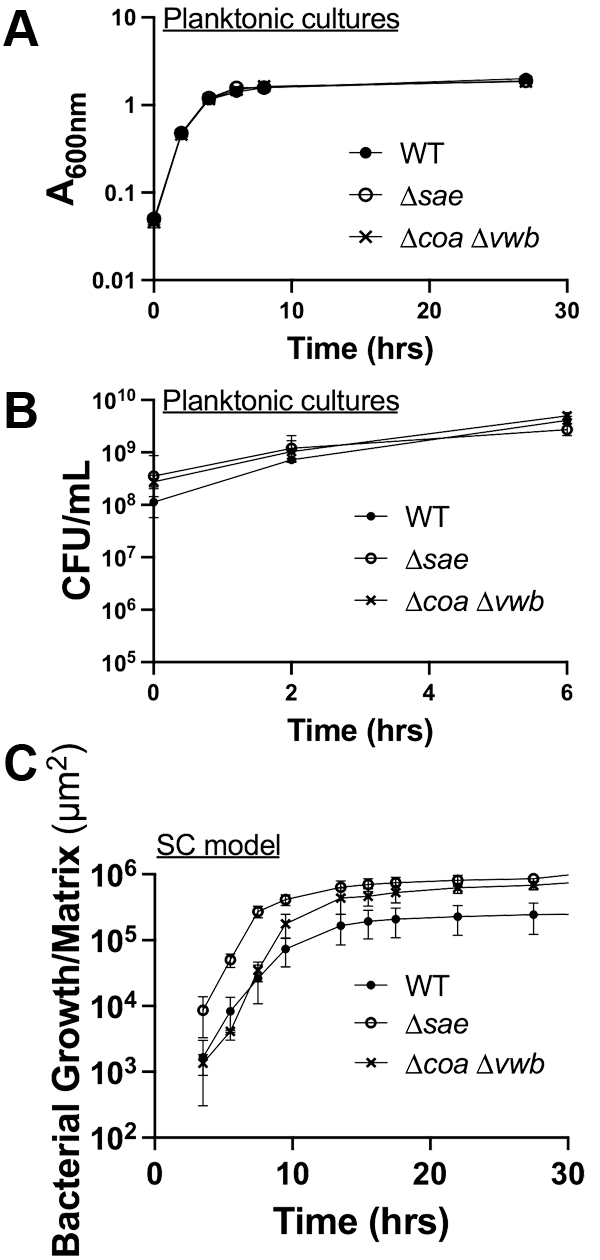

### Supplemental Fig 3

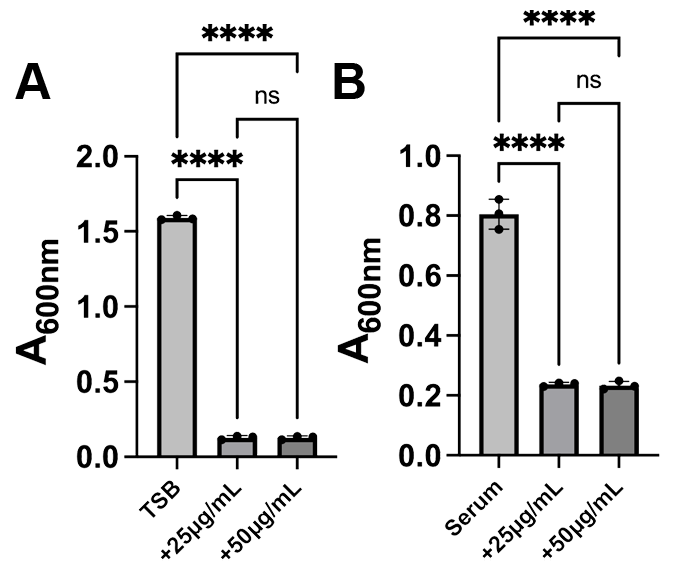
